# Sensitization with an allogeneic MHC class I molecule induces anti-tumor immunity in the absence of PD-1 in mice

**DOI:** 10.1101/2023.08.26.554968

**Authors:** Komang Alit Paramitasari, Yasumasa Ishida

## Abstract

To investigate the effect of a major histocompatibility complex class I (MHC-I) overexpression to augment immune sensitivity against tumors, we have generated the murine colorectal carcinoma cell line MC38 (with the endogenous H-2^b^ haplotype) overexpressing the allogeneic mouse MHC-I cell surface molecule H-2K^d^ (MC38 H-2K^d^). The tumorigenicity of unmodified parental cells (MC38 PT) and MC38 H-2K^d^ was tested *in vivo* by subcutaneous injection into the flank of wild-type (WT) and programmed death-1 (PD-1) knockout (KO) mice in a C57BL/6 (H-2^b^) genetic background. MC38 PT cells readily formed tumors and grew progressively in both WT and PD-1 KO mice. The speed of MC38 PT tumor growth was slower in PD-1 KO mice than in WT mice. In contrast, MC38 H-2K^d^ cells showed full sensitivity to rejection by the immune system in both naïve WT and PD-1 KO mice, indicated by spontaneous tumor regression. Next, we sought to determine the extent to which H-2K^d^-overexpressing tumors could protect the mice against unmodified cancers. PD-1 KO mice were first sensitized with highly immunogenic MC38 H-2K^d^ cells and then challenged with weakly immunogenic MC38 PT cells. Intriguingly, all PD-1 KO mice gained immunity against the aggressive MC38 tumor and became tumor-free. Sensitizing PD-1 KO mice with growth-arrested (by the pre-treatment with mitomycin C, MMC) and the debris of MC38 H-2K^d^ tumors also provided full protection against the growth of secondary MC38 PT tumors. Most notably, sensitization with the debris of MC38 H-2K^d^ cells provided the long-term immunological memory against MC38 PT carcinoma cells. This finding implies that MC38 H-2K^d^ cells retain highly efficient and durable immunogenicity.

## **1.** Introduction

### Programmed death-1 (PD-1) as a negative regulator of immune responses

PD-1 is a type-I transmembrane protein, and its expression is strongly elevated upon antigenic stimulation of T cells. The PD-1 gene was discovered in 1991 by exploiting the subtractive-hybridization technique (1). Several years later, observation of PD-1 deficient mice revealed that PD-1 negatively regulates immune responses (2). PD-1 has two ligands, namely programmed death-ligand 1 (PD-L1, also called B7-H1) and PD-L2 (B7-DC). The PD-L1 molecule is expressed on a wide variety of hematopoietic and non-hematopoietic cell types, whereas the expression of PD-L2 is restricted to professional antigen-presenting cells (APCs), in which dendritic cells (DCs) and macrophages are included. The PD-1 cytoplasmic region has two tyrosine residues in the immunoreceptor tyrosine-based inhibitory motif (ITIM) and the immunoreceptor tyrosine-based switch motif (ITSM), which is essential for the negative-regulatory activity of PD-1. Upon engagement of PD-1 with its ligand, the tyrosine residue in ITSM is phosphorylated and further recruits the SHP-2 tyrosine phosphatase. Consequently, the SHP-2 activity on PD-1 suppresses the TCR signaling and T cell-mediated immune responses (3–6).

### Cancer immunotherapy with PD-1 checkpoint blockade

In recent years, the PD-1–PD-L1 pathway has shed some light on the improvement of antitumor immunity (7,8). PD-1 is expressed on activated T cells and further dampens down the cytotoxic T lymphocyte (CTL) activity upon ligation with PD-L1 expressed on tumor cells. By inhibiting the PD-1/PD-L1 pathway through the administration of the anti-PD-1 (or anti-PD-L1) monoclonal antibody, the T cell could be reactivated and gain cytotoxic activity against tumor cells. This finding led to the first clinical study of anti-PD-1 antibodies in 2006, which was associated with antitumor immunity for several solid tumors (9). Furthermore, the therapeutic antitumor immunity targeting the PD-1/PD-L1 checkpoint pathway has been attributed to durable response towards multiple types of cancers including non-small cell lung cancer (NSLC), renal cell carcinoma (RCC), bladder carcinoma, melanoma, and Hodgkin’s lymphoma (10–21).

The prominent results from clinical studies of PD-1 blockade have attracted scientists’ interest to exploit the PD-1 molecules in the targeted therapies. Cancer immunotherapy mediated by PD-1/PD-L1 checkpoint inhibition (notably that using nivolumab or pembrolizumab) has received much attention in recent years due to its effectiveness to eliminate tumors. Treatment using such fully human anti-PD-1 antibodies has been demonstrated significant outcomes with durable, long-lasting responses (22), even in patients whose immunotherapy has been discontinued. However, major subsets of patients are refractory to the benefits of PD-1 therapy. Hence, the responses to the PD-1 blockade still vary among different types of cancer cells and patients. One of the major obstacles in cancer immunotherapy is the heterogeneity of tumors. Not all tumors appear to respond to cancer immunotherapy (9, 23, 24). These findings show significant implications that particular cancer may exhibit immune escape mechanisms.

The most common tumor escape mechanism is demonstrated in tumors with a lack of antigenic expression and loss of MHC-I proteins where they cannot be recognized by T cells and become invisible to the immune system. In parallel, cancer cells can exploit distinct immune evasion mechanisms by opting the PD-1/PD-L1 inhibitory pathway. It has been described that tumors with poor immunogenicity acquired resistance to PD-1 blockade due to defects in antigen-presenting machinery, failing the recognition of tumor antigens (25–27).

### MHC-I as the whistleblower of anti-tumor immune response

Class I MHC molecules are constitutively expressed on almost all nucleated cells. Biological structure of MHC-I molecules is composed of a polymorphic α chain (or heavy chain) non-covalently linked to the β2-microglobulin. The α1 and α2 domains contain highly polymorphic residues of amino acids where they function to provide variations in peptide binding and T-cell recognition. Accordingly, the expression of MHC-I molecules provides an important display system for tumor antigens (28–31).

It is generally accepted that the presentation of tumor antigens by MHC-I is required for the successful PD-1/PD-L1 blockade. In support of this point, initial attempts focusing on the induction of MHC-I expression have been shown potential outcomes to enhance the tumor-specific immune response (32–34). In addition, the transfer of an allogeneic mouse MHC-I gene was proven to significantly enhance the immunogenic property of resistant cancer cells. Immunization of mice with melanoma cells stably transfected with a cytokine gene alone, or particularly in combination with an allogeneic mouse MHC-I gene, was reported to stimulate CTL responses and attain a higher survival rate in immunized mice (35).

It is well established that the MHC alloantigens provoke very strong CTL-restricted immune responses, thus resulting in rapid graft rejection after transplantation (36). This suggests that the recipient T cells are capable of recognizing the peptide-allogeneic MHC complexes, an immune mechanism well known as allorecognition or alloreactivity. Recognition of alloantigens by alloreactive T cells falls into two categories: direct and indirect recognition (37). In the direct pathway, the intact MHC molecules are recognized through TCRs of recipient T cells. Despite the conventional allorecognition of peptide-MHC complexes by TCR, some models propose that the TCR may interact either with the peptide portions or the MHC molecules alone (38). This implies that both MHC and peptides play central roles in activating T cells. On the other hand, indirect recognition presumed the allogeneic MHC molecules are engulfed and processed by the recipient antigen-presenting cells (APC). Hence, the resulting peptides derived from allogeneic MHC are bound and presented by the host (self) MHC for the further cross-priming of T cells. This leads to the conventional immune response towards foreign antigens. Moreover, the engulfment of allogeneic MHC molecules through endocytic vesicles may end up in the MHC class II pathway that would also lead to the allorecognition by CD4^+^ T cells. It has been confirmed that the CD4^+^ T-cell responses are also beneficial for tumor elimination (39–45).

### Cancer therapy in combination with PD-1 checkpoint blockade

A myriad of reports have shown that the combination therapies with the PD-1 blockade are more effective than a single treatment (46–48). However, little evidence is available to elucidate the function of allogeneic MHC-I in overcoming resistance to PD-1 therapy. Indeed, the study to address the effect of the PD-1/PD-L1 checkpoint inhibition in combination with the enhancement of MHC-I expression has become urgent to be accomplished. Interestingly, the combination of radiotherapy with immune checkpoint blockade has demonstrated a synergistic effect to improve the abscopal effect, a rare phenomenon of tumor regression that is located not only at the primary irradiated sites but also in the non-irradiated, distant sites (49–55). Investigation of the abscopal effect as the results of PD-1 therapy combined with other immune adjuvants has gained attention in the immuno-oncology field.

The objective of this study is to address a possible experimental system for the enhancement of tumor immunity in the context of PD-1/PD-L1 blockade. Tumor cells may develop immune evasion mechanisms due to failure in the antigen-presenting machinery. The introduction of exogenous MHC-I in combination with PD-1 immune checkpoint blockade may augment the tumor-antigen recognition and thereby provide antitumor responses. In the present study, an expression vector harboring one of the mouse MHC-I genes (H-2Ks), namely allogeneic H-2K^d^, was constructed. Moreover, the established construct was then stably introduced into murine cancer cells with a C57BL/6 genetic background (H-2^b^) in the presence (wild-type [WT] mice) or absence of the PD-1 gene (PD-1 knockout [KO] mice). The modified cancer cells were harnessed in order to investigate the immune reaction towards alloantigens on the basis of PD-1 checkpoint inhibition. Our hypothesis is that combination therapy will augment the tumor immune response and induce an abscopal effect, resulting in a robust global anti-tumor response.

## 2. Materials and Methods

### Generation of the MHC-I expression vector

The H-2K^d^ cDNA was amplified by reverse transcriptase-mediated polymerase chain reactions (RT-PCR) using total RNA extracted from the spleen of BALB/c mice and cloned into the pCAGGS plasmid (56). After the cloning, the nucleotide sequence of the H-2K^d^ cDNA was verified by the standard sequencing procedures.

### Tumor cell lines

MC38 (57) and MCA-205 (58) cell lines were cultured in high-glucose Dulbecco’s-modified Eagle’s medium (Sigma Aldrich) supplemented with 10 % (v/v) of heat-inactivated fetal bovine serum (FBS), 2 mM L-glutamine (Gibco), and antibiotics (100 µg/ml streptomycin and 100 U/ml penicillin, Gibco). Cells were grown inside a 37 °C tissue culture incubator maintained at 5% CO_2_. All cells used in this study were tested free for mycoplasma contamination, confirmed with e-Myco mycoplasma PCR detection kit ver.2.0 (LiliF Diagnostic) according to the manufacture’s protocols.

### Stable transfection

One day prior to transfection, 5 × 10^4^ MC38 or MCA-205 cells were seeded on a 24-well dish in the growth medium without the presence of antibiotics. Transfections were performed in a total volume of 550 µl of growth medium containing 3 µg/µl of Polyethylenimine (PEI-Max) and a mixture of 1 µg of *Sal*I-linearized plasmid DNA encoding mouse allogeneic MHC-I genes and 100 ng of *Sca*I-linearized KT3NP4 plasmid (59) harboring neomycin-resistant gene (Neo^R^). After 24 h, transfected cells were replated in 10 ml of complete growth medium containing 500 µg/ml of G418 (Sigma Aldrich) as the selection medium. The G418-containing medium was replaced every 2 days to maintain the selective pressure. After 10-12 days, the single-isolated Neo^R^ colonies were harvested using stainless steel cloning cylinder sealed with sterile silicone grease and transferred into a 24-well dish. Each colony was screened through flow cytometric analysis to detect the expression of transgenes.

### Flow cytometry (FACS) analysis of stable clones

Once satisfactory confluency (80-90 %) of isolated cells in a 24-well dish was reached, the old-growth medium was discarded. Each well was rinsed twice with 1X PBS and 100 µl of 0.25 % Trypsin-EDTA (Gibco) was added into the cells. When the cells have begun to round up and detached from the dish bottom, 400 µl of growth medium was added to quench the trypsin. Each cell in the well was transferred into 1.5 ml tubes and spun down at 1,000 x *g* for 5 minutes at 4 °C. The supernatant was aspirated without disturbing the cell pellet. The centrifugation and aspiration steps were repeated twice to ensure the complete removal of growth medium traces. In the final washing step, the cells were suspended in a 100 µl of FACS staining buffer (1 % FBS and 10 mM EDTA in 1X PBS). All cells were stained for 30 min on ice with the indicated fluorochrome-conjugated antibodies; anti-H-2K^d^ PE (clone SF1-1.1.1, eBioscience), anti-H-2K^b^ APC (clone AF6-88.5.5.3, eBioscience), biotinylated. Stained cells were detected by BD Accuri C6 flow cytometer, and the data were acquired using CFlow software.

### Limiting dilution cloning

For the limiting dilution cloning, an average density of 0.5 cells/well of stably transfected MC38 or MCA-205 tumor cells were cultured in a 96-well dish. Cells were grown in a conditioned medium (0.2 µm sterile-filtered old medium derived from cells 24 h post-culture). The cells were incubated (37 °C, 5 % CO_2_) for 14-21 days until the colonies were observed. Colonies derived from single cells representing a monoclonal population were harvested and cultured in a 24-well dish. Isolated monoclonal cells were subjected to flow cytometry to screen the allogeneic MHC-I surface expression. Clones that expressed the highest allogeneic MHC-I were expanded in a 10-cm dish and cryopreserved for the sequential *in vivo* experiments.

### Animals

All experiments were conducted in accordance with the animal experimentation guidelines of the Nara Institute of Science and Technology that comply with the National Institute of Health guide for the care and use of laboratory animals. All efforts were made to minimize the number of animals used and their suffering. PD-1 KO mice (2) and WT mice (CREA Japan) in a C57BL/6N genetic background were bred and maintained in a specific pathogen-free environment and climate/light cycle controlled animal room at a constant temperature of 23-24 °C, 50-70 % humidity with *ad libitum* access to pellet diet and water. Female BALB/c ^nu/nu^ immunodeficient mice (CLEA Japan) were used between 8-9 weeks of age. Humane endpoints were applied to prevent unalleviated pain according to guidelines. Tumor volume was measured with digital calipers and animals were euthanized before tumor size reached the maximum allowed or if the tumor displayed a sign of ulceration. Euthanasia was performed according to guidelines for animal cervical dislocation.

### Allogeneic tumor studies

*In vivo* tumor studies were performed as follows: age-matched 8-12 week-old male/female PD-1 KO and/or WT mice were inoculated subcutaneously in the unilateral or bilateral flank with allogeneic MHC-I-expressing tumor cells. Tumor injections were prepared at 5×10^6^ cells/ml concentration suspended in Dulbecco’s PBS (-) (Nacalai Tesque) inside a 1ml syringe with a 27G needle (Terumo). Mice were anesthetized using continuous isoflurane gas followed by subcutaneous inoculation of cancer cells in the dorsal flank region. Each tumor injection had a volume of 100 µl containing 5×10^5^ cells. Palpable tumors were measured starting day 5-6 post-inoculation and every 2 days thereafter until the endpoint of the experiment. Tumor volume was calculated using the formula: 0.5 x length x width^2^, where the length represents the largest tumor diameter and width represents the perpendicular tumor diameter.

### Bilateral tumor model

Bilateral tumor studies were carried out with the objective to achieve the abscopal effect. The same number of unmodified or parental (PT) and H-2K^d^ cells were administered subcutaneously in both flanks.

### Sensitization with live cells

A total of 5×10^5^ cells expressing MC38 H-2K^d^ in 100 µl of D-PBS (-) suspension were inoculated into the left flank of PD-1 KO mice. Mice were challenged with 5×10^5^ parental cells in the contralateral flank after six days.

### Sensitization with growth-arrested cells

The MC38 H-2K^d^ stable clones were seeded one day prior to mitomycin C (Fujifilm Wako) treatment and grown in humidified culture incubator until they reached 70% confluency on the next day. A total concentration of 10 µg/ml of mitomycin C (MMC) reconstituted in sterile D-PBS (-) was administered into the culture medium and the cells were incubated for 3 h. The culture medium containing MMC was replaced with pre-warmed complete medium after washing the cells with 1X PBS two times. The growth-arrested cells were harvested on the next day for s.c. injection (5×10^5^) into the left flank of mice. On day 6 the mice were given the same dose of injection of parental unmodified cells in the right flank.

### Sensitization with cell debris

MC38 H-2K^d^-harboring cells were aseptically collected and adjusted to reach the concentration of 5×10^6^ cells in 1 ml of D-PBS (-) and then transferred into cryogenic vials. The vials were submerged into liquid nitrogen for 5 minutes and immediately transferred into prewarmed water at a temperature of 37 °C until the entire cells were completely thawed. These freeze-thaw steps were repeated 5 times and then mixed vigorously to prevent cell aggregation. Complete cell death was assessed by trypan blue exclusion. Cell debris was injected into the left flank and marked as day 0. On day 6, the same mice were implanted with 5×10^5^ parental cells in the right flank.

### Criss-cross tumor model

To examine the specific immunity between MC38 and MCA-205 tumors, the criss-cross experiment was performed in which the mice were subsequently challenged with secondary tumors disparate from the pre-established parental tumors. WT mice were given 5×10^5^ of MC38 H-2K^d^ tumor cells as a primary tumor in the left flank. After six days, the MCA-205 PT cells were implanted in the right flank as the secondary tumor. On the other hand, 6 days after the establishment of s.c. MCA-205 H-2K^d^ tumors, PD-1 KO mice were challenged with MC38 PT in the opposite flank.

### Assessment of tumor-infiltrating lymphocytes (TILs)

MC38 or MCA-205 tumors were resected from euthanized tumor-bearing mice. Bulk tumors were prepared by cutting tumors with scalpel blades, followed by enzymatic digestion with 0.13 Wünsch units of Liberase TL (Roche) in 10 ml of RPMI-1640 (Fujifilm Wako) for 45 minutes at 37 °C with gentle agitation (30 r/min). The digested tumor was passed through a 40 µm cells strainer (Falcon) and washed with 1X PBS (containing 2 % FBS and 10 mM EDTA to stop the enzymatic reaction) and spun down at 1,000 x *g* for 10 min at 4 °C. The tumor was undergone density separation using a discontinuous isotonic Percoll gradient (GE Healthcare Life Sciences) to remove debris and necrotic cells. One part of 1.5 M NaCl was added to 9 parts of Percoll (1.130 g/ml) to create 100 % Percoll solution, which was then diluted to 80 %, 40 %, and 20 % solutions with 0.15 M NaCl. The filtered cell suspension was washed and suspended in 4 ml of 80 % Percoll, which was subsequently overlaid with 4 ml of 40 % Percoll, and then 3 ml of 20 % Percoll above it. Tubes were centrifuged at 1,000 x *g* for 30 min at room temperature. The resulting interfacial layer between the 80 % and 40 % layers was collected for flow cytometric analysis.

### Flow cytometry of lymphocytes

Splenocytes were extracted by removing the spleen aseptically and crushed with cover glasses. The cell suspension was passed through 40 µm nylon mesh and subjected to erythrocytes lysis by using HybriMax red blood cell lysis buffer (Sigma Aldrich) at room temperature for 3 minutes. The cells were washed twice with 1X PBS and spun down at 1,000 x *g* for 3 min at 4 °C. Inguinal and axillary lymph nodes on both sides of the mouse body were collected and crushed in the suspension of FACS Staining Buffer (1 % heat-activated FBS and 10 mM EDTA in 1X PBS) and filtered through 40 µm nylon mesh. The cells were washed twice with FACS Staining buffer and centrifuged at 1,000 x *g* for 3 min at 4 °C. Isolated lymphocytes derived from spleen, lymph nodes, and tumors were incubated with purified rat anti-mouse CD16/CD32 (FCɣIII/II Receptor, clone 2.4G2, BD Pharmingen) for 10 min and directly stained with APC-conjugated anti-mouse CD45 (clone 30-F11, BD Pharmingen) to specifically stained leukocytes. Cells were stained for other surface markers: CD3ε (clone 145-2C11, eBioscience), CD4 (clone GK1.5, eBioscience), CD8α (clone 53-6.7, eBioscience), PD-1 (clone RMP1-30, eBioscience), and propidium iodide (eBioscience). Cells were analyzed on the BD Accuri C6, and data were processed using CFlow software.

### Matrigel-embedded cancer cell study

PD-1 KO mice were subcutaneously injected with a mixture of MC38 cells and growth factor-reduced Matrigel matrix (phenol red-free) (Corning). A total of 5×10^5^ MC38 parental or H-2K^d^-transfected cells (live or debris) was suspended in 700 µl of ice-cold Matrigel and kept on ice until injection. All equipments (syringe, tips, and needles) and reagents were chilled at all times to prevent the polymerization of Matrigel matrix. Before injection, mice were anesthetized with continuous isoflurane gas. The cells-Matrigel mixtures were loaded into a chilled syringe and the needle was injected all the way to the end in order to reduce leaking. The syringe was held in place for at least 30 second to allow the gelling to begin. A bump will appear at the injection site (the size will reduce due to absorption and partial degradation of Matrigel *in vivo*).

### Hematoxylin and eosin (HE) staining

At the endpoint of the Matrigel-related experiment (day 7), the Matrigel plugs were removed by wide excision of the flank subcutaneous tissue. Each plug was embedded in Tissue-Tek OCT compound (Sakura), immediately snap-frozen on liquid nitrogen vapors, and cut into 8-µm sections using a cryostat. Frozen tissue sections were air-dried overnight and fixed with acetone for 5 minutes at room temperature. All of the following steps for HE staining were performed at room temperature. Cryosections were rehydrated in distilled water for 5 minutes and nuclei were stained with Mayer’s hematoxylin solution (Fujifilm Wako) for 5 minutes and then rinsed with running tap water. Sections were stained with eosin and dehydrated consecutively in 80 %, 90 %, and absolute ethanol, then further bathed with xylene three times before mounting with PathoMount solution (Fujifilm Wako).

### Immunohistochemistry

For immunostaining, upon acetone fixation, sections were incubated for 1 hour at RT with D-PBS (-) + 5 % normal goat serum (Abcam) to block nonspecific binding sites and then incubated overnight at 4 °C with primary antibodies diluted in D-PBS (-). Following primary antibody staining, sections were washed three times with 1X PBS and incubated with fluorochrome-conjugated secondary antibodies diluted in D-PBS (-) for 1 hour at RT. Secondary antibodies were then washed with 1X PBS three times and the nuclei were stained with DAPI (4′,6-diamidino-2-phenylindole). Slides were finally mounted with Immunoselect antifading mounting medium (DNV Dianova). Immunostaining was performed using the following primary antibodies: rat anti-mouse CD8α (clone 53-6.7, eBioscience) and Armenian hamster anti-mouse CD11c (clone N418, BioLegend). CD8 primary staining signal was further amplified using an anti-rat IgG secondary antibody conjugated with Alexa Fluor (AF) 555. An anti-Armenian hamster IgG antibody conjugated with AF647 was used in combination with CD11c primary antibody. Samples were imaged using a confocal laser scanning microscope LSM 710 (Carl Zeiss) and images were processed using ImageJ (National Institute of Health).

### Statistical analysis

Statistical analysis was performed using Prism 9.0 (GraphPad Software). Differences between 2 groups were evaluated using unpaired Student’s *t*-test with Welch’s correction unless otherwise stated. Kaplan-Meier survival curves were analyzed with log-rank (Mantel-Cox) test. The level of significance was set at \**p* < 0.05, \*\**p* < 0.01, and *** *p* < 0.001.

## 3. Results

### Transfer of the allogeneic H-2K^d^ gene to MC38 and MCA-205 cancer cells

We introduced the expression vector for the allogeneic H-2K^d^ molecule into MC38 and MCA-205 cancer cells. To determine the expression of syngeneic and allogeneic class I MHC molecules from the parental and transfected cells, FACS analysis was employed using anti-mouse H-2K^b^ and H-2K^d^ monoclonal antibodies. We confirmed that MC38 and MCA-205 cells constitutively expressed high basal levels of endogenous H-2K^b^ class I MHC (Fig. 1 and Fig. 2). In the *in vitro* study, the stably transfected MC38 and MCA-205 cells successfully overexpressed high levels of allogeneic H-2K^d^. We therefore concluded that the transfectants became double-positive for both H-2K^b^ and H-2K^d^ expressions (Fig. 1 and Fig. 2). The expression of H-2K^d^ was stable after more than 5 months of continuous passage in culture.

**Figure 1.**
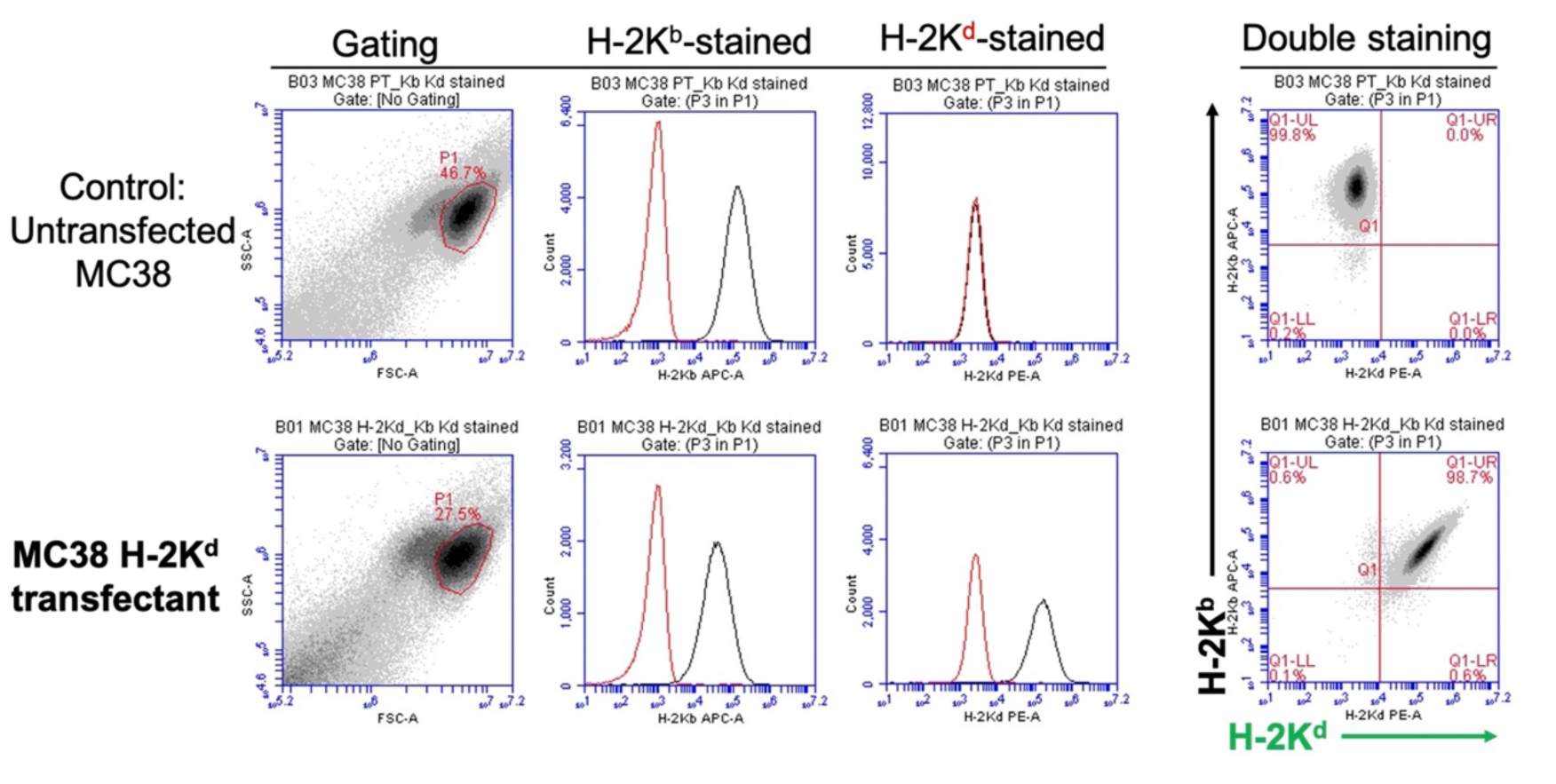
Flow cytometric assessment of unmodified and stably transfected MC38 cells. The transfectant simultaneously expressed H-2K^d^ and H-2K^b^ class I MHC molecules represented by the entire shifting of histogram plot into the right side (black line) and position of the cell population into the upper right quadrant in the density plot. Unstained control is shown in the red line. Data are representative of at least five independent experiments.

**Figure 2.**
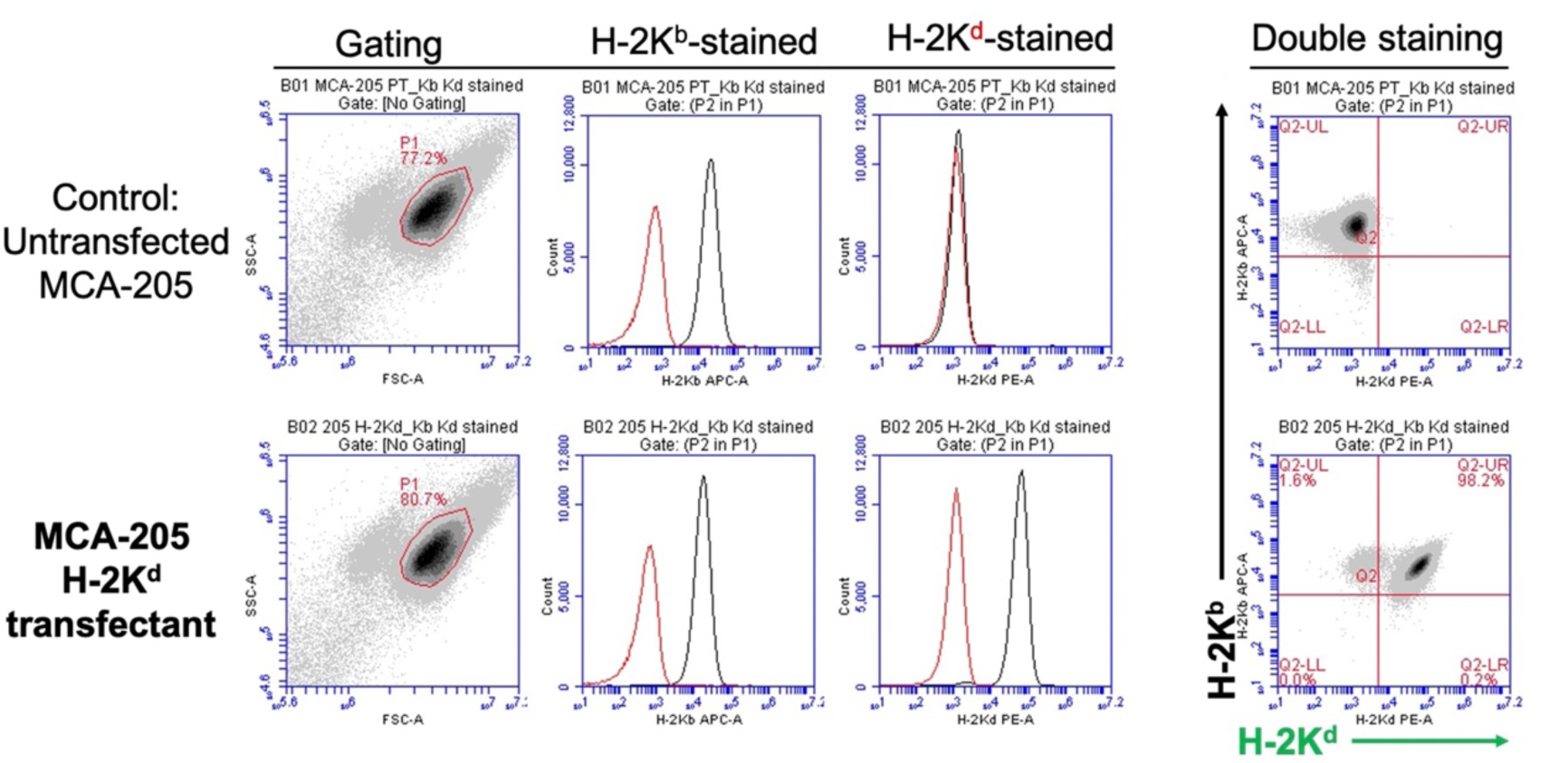
Flow cytometric assessment of unmodified and stably transfected MCA-205 cells. The transfectant simultaneously expressed H-2K^d^ and H-2K^b^ class I MHC molecules represented by the entire shifting of histogram plot into the right side (black line) and position of the cell population into the upper right quadrant in the density plot. Unstained control is shown in the red line. Data are representative of at least five independent experiments.

### Interferon-gamma (IFNɣ) stimulation profoundly elevates the cell-surface expression of PD-L1 and MHC-I

Next, we asked whether the MC38 and MCA-205 cancer cells innately expressed immunoregulatory ligand PD-L1. The basal (unstimulated) level of PD-L1 in MC38 cells was moderate while MCA-205 displayed a low level of PD-L1 expression (Fig. 3 and Fig. 4). High PD-L1 inhibitory ligand levels are usually associated with poor prognosis (60), and confirmation of enhanced PD-L1 expression *in vitro* upon cytokine stimulation might be indicative for the actual condition in the tumor microenvironment where IFNɣ is secreted by the effector T cells.

**Figure 3.**
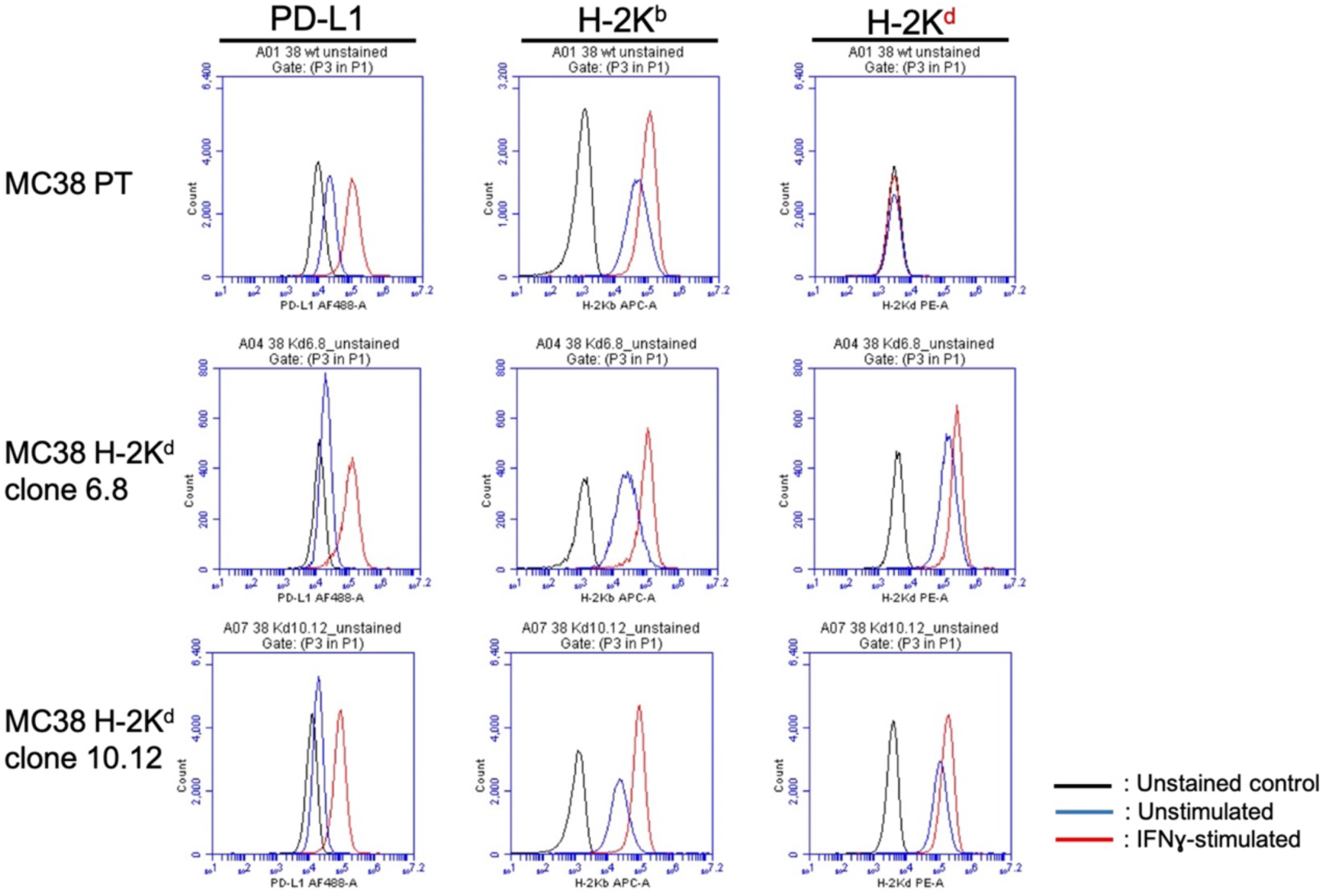
Transfected MC38 clones (see the main text for Fig. 7 for details) were cultured *in vitro* and stimulated with IFN-γ (20 ng/ml) for 24 h. Expression of PD-L1, H-2K^b,^ and H-2K^d^ was assessed by flow cytometry. Unstained control is shown in the black line while unstimulated cells are represented in the blue line. Data are representative of two independent experiments.

**Figure 4.**
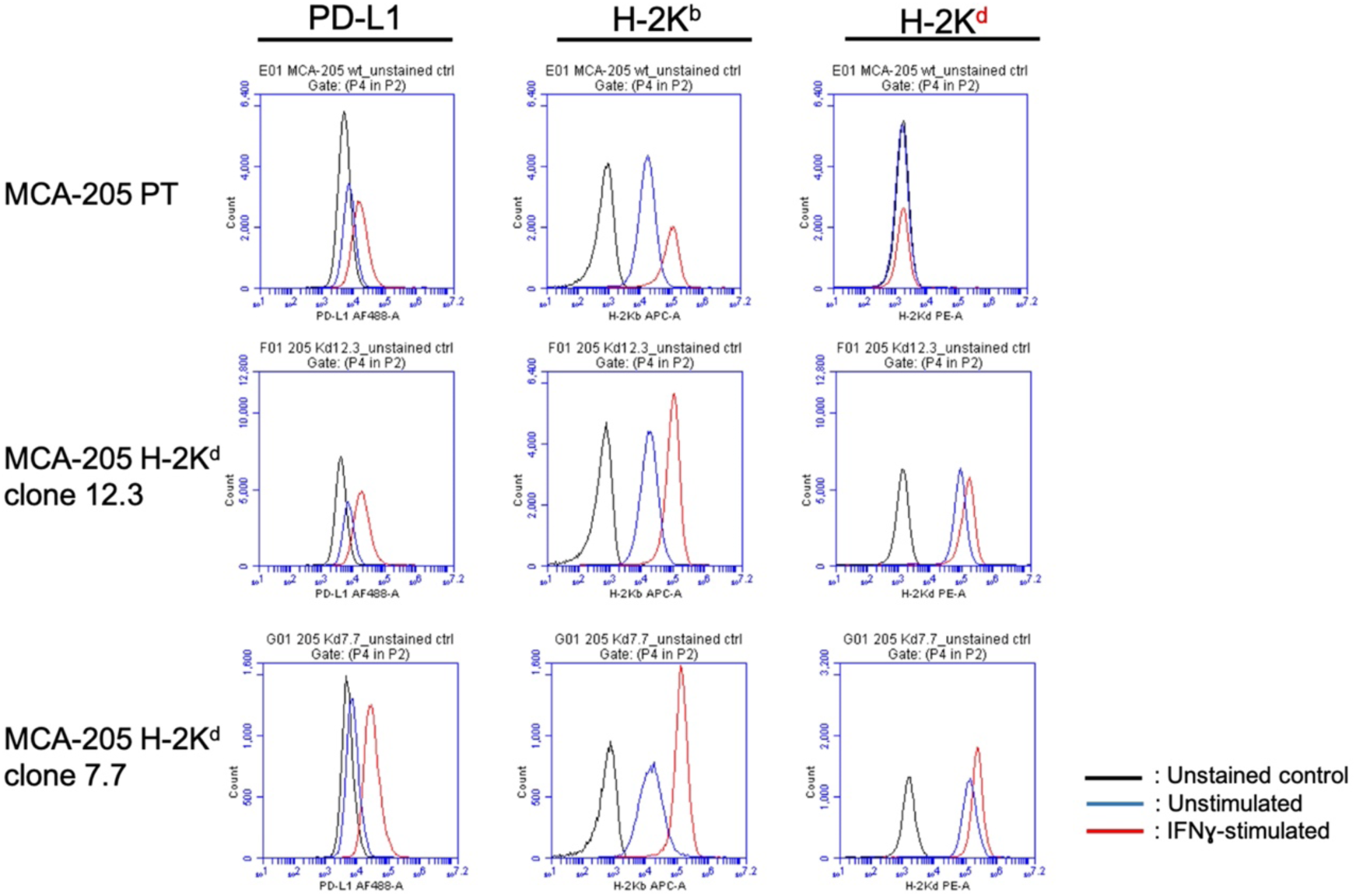
Transfected MCA-205 clones (see the main text for Fig. 9 for details) were cultured *in vitro* and stimulated with IFN-γ (20 ng/ml) for 24 h. Expression of PD-L1, H-2K^b^ and H-2K^d^ was assessed by flow cytometry. Unstained control is shown in the black line while unstimulated cells are represented in the blue line. Data are representative of two independent experiments.

Therefore, the original and H-2K^d^-overexpressing stable clones were stimulated with 20 ng/ml IFNɣ for 24 h (60) to assess the expression of PD-L1. Although the mechanism as to how the IFNɣ can upregulate MHC-I and PD-L1 expression has not been elucidated in detail, it was reported that IFNɣ affected post-translational regulation of genes involved in antigen-presenting machinery (61).

IFNɣ treatment evidently boosted PD-L1 expression, as well as endogenous and exogenous MHC-I (H-2K^b^ and H-2K^d^, respectively) in MC38 (Fig. 3) and MCA-205 cells (Fig. 4). Inducibility of the PD-L1 expression in MC38 cancer cells was higher than that of MCA-205. These results might be considered to predict the possibility of PD-1/PD-L1–mediated antitumor immune resistance in the *in vivo* settings. The two-faced effects of IFNɣ presence could be beneficial to increase MHC-I expression, while on the hostile side, could also induce inhibitory PD-1-receptor signaling.

### MC38 and MCA-205 tumors show different levels of sensitivity to PD-1 deficiency

MC38 colon adenocarcinoma cells have been previously reported as an immunogenic cancer cell line with high sensitivity to PD-1/PD-L1 axis blockade (60, 62–67). In this study, MC38 cells appeared to be partially responsive to the global absence of PD-1 in the KO mouse model studied. Subcutaneously implanted parental MC38 cells (MC38 PT) formed tumors and grew progressively in both PD-1 KO and WT mice (Fig. 5A), but the speed of tumor growth appeared to be slower in PD-1 KO mice (Fig. 5B). Robust tumor advancement followed by ulceration within two weeks post-inoculation resulted in low survivability of tumor-bearing mice.

**Figure 5.**
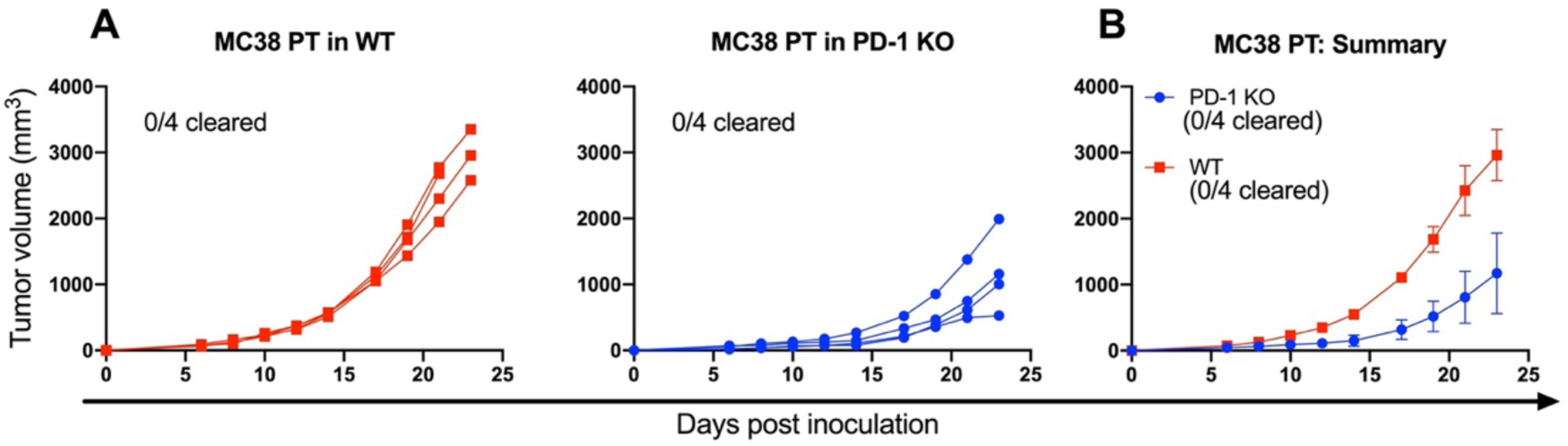
Subcutaneous injection of parental (unmodified) MC38 cells into WT and PD-1 KO mice resulted in progressive tumor growth. Individual tumor growth curves **(A)** and average tumor growth curves **(B)** are shown. Data are representative of at least two independent study repeats with n = 4 to 5 mice per group. Error bars depict the standard error of the mean (SEM) from the means.

In contrast, the methylcholanthrene (MCA)-induced murine sarcoma cell line MCA-205 was shown to be highly responsive to PD-1 deficiency (Fig. 6A and 6B). The MCA-205 tumor implant was initially formed in PD-1 KO mice and then completely cleared. Tumor recurrence has never occurred for at least for six months of observation and the animals became tumor-free. In contrast, no tumor regression was seen in WT mice. In this study, the unleashed antitumor immune responses supported by deficiency of PD-1 inhibitory pathway were detrimental for MCA-205, but not for MC38 cancers.

**Figure 6.**
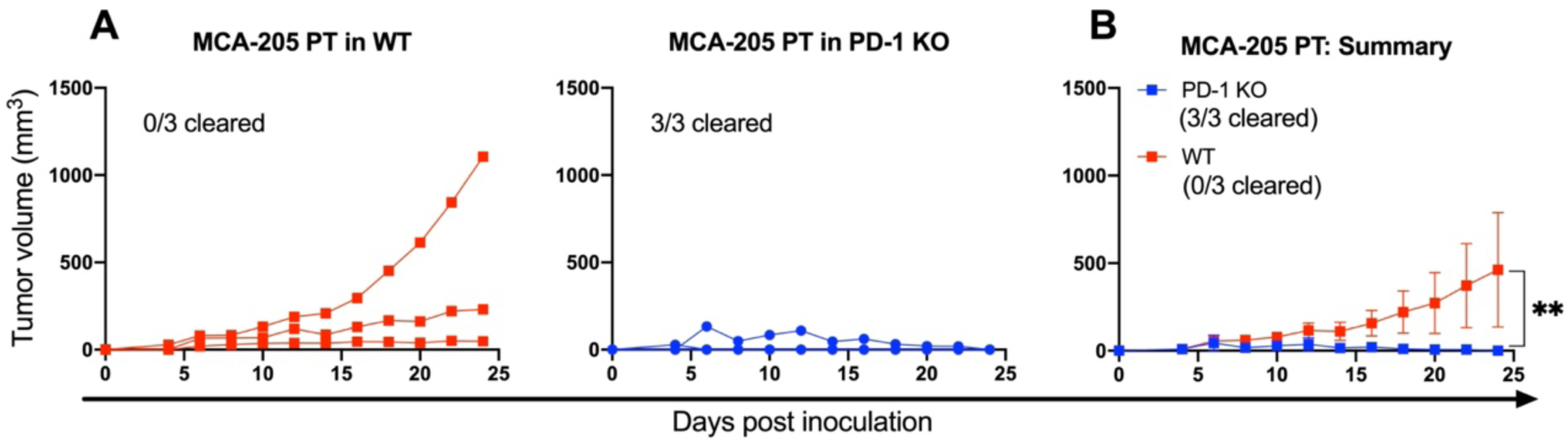
MCA-205 PT cells were inoculated subcutaneously into the flank of PD-1 KO and WT mice. **(A)** Individual tumor growth curves in PD-1 KO mice showed complete regression (blue lines) while in WT mice, tumors were growing progressively (red lines). **(B)** Data of the tumor growth curves are summarized. Representative results from one of three repeated experiments with n = 3 mice per group are shown. Statistical significance was determined by unpaired Student’s *t*-test. Error bars depict means ± SEM.

These observations demonstrated a likelihood of immune evasion exploited by MC38 and MCA-205 tumors. Despite the apparent fact that both tumors were competent at expressing their own endogenous H-2K^b^ molecules *in vitro*, the implanted tumors nevertheless escaped from immune surveillance. As previously reported, on the other hand, a group of cancer cells could evade the recognition of CD8 T cells by downregulating the MHC-I antigen presentation pathway through gene mutation and/or epigenetic modification (68).

### Allogeneic H-2K^d^ overexpression in MC38 provides complete tumor regression in WT and PD-1 KO mice

To validate that allogeneic MHC-I was attributable specifically to enhance host immunity against tumors, each MC38 H-2K^d^ bulk line was subjected to the limiting-dilution experiment, and single-cell clones were then inoculated into the mice. Indeed, the monoclonal MC38 H-2K^d^ cells (clone #6.8) were completely regressed in PD-1 KO and WT mice (Fig. 7A), signifying high immunogenicity. Next, to exclude the possibility that H-2K^d^-mediated tumor regression might be effective only to a particular clone, we inoculated a different clone (MC38 H-2K^d^ #10.12) and confirmed the growth and spontaneous regression in both WT and PD-1 KO mice (Fig. 7B). To this end, the clonal variability was not affecting the *in vivo* study. Taken together, these findings undeniably suggested that allogeneic H-2K^d^ involvement is required for successful tumor clearance. In addition, the regression of the MC38 H-2K^d^ tumor was confirmed to be dependent on allogeneic MHC-I itself given that the control of MC38 tumor encoding Neo^R^ gene alone grew progressively (Fig. 7C).

**Figure 7.**
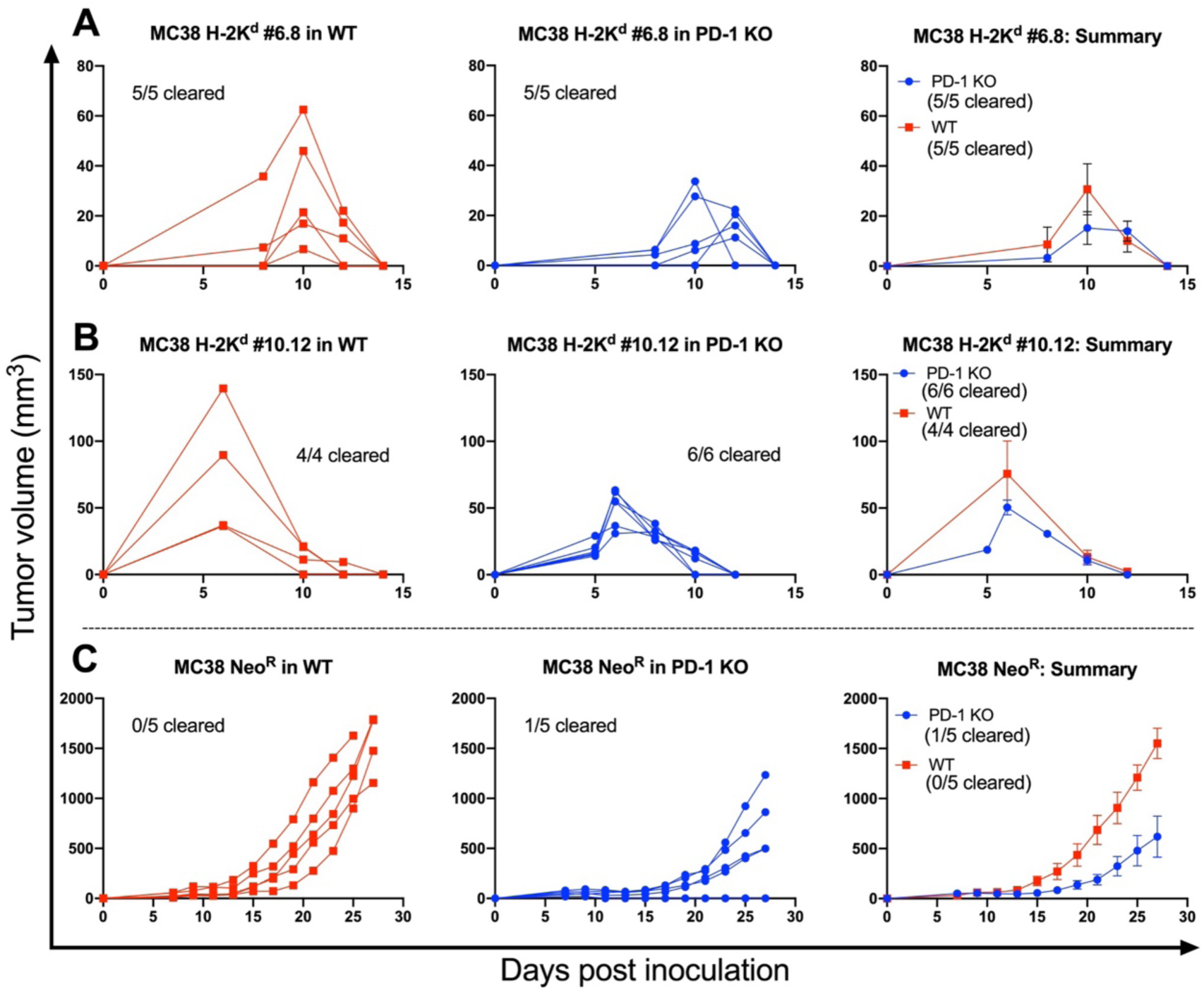
MC38 tumors expressing allogeneic H-2K^d^ were completely rejected from the PD-1 KO and WT mice. Tumor growth curves of MC38 H-2K^d^ from clones #6.8 **(A)** and #10.12 **(B)** are shown. **(C)** MC38 Neo^R^ tumors were used as controls to exclude the possibility of drug resistance-mediated immunogenicity. Data shown are from two independent experiments with n = 5 mice per group (A) and from one experiment with n = 4-6 mice (B and C). Average tumor growth curves depict means ± SEM.

Having observed a tremendous immune response demonstrated by the expression of allogeneic H-2K^d^ in MC38 tumors, we further asked whether this result was specifically due to H-2K^d^–mediated modulation of antitumor immunity rather than the possibility of delayed tumor onset. To clarify this point, we compared the tumorigenicity of MC38 cells in an immunodeficient mouse model (Fig. 8A). BALB/c^nu/nu^ mice lack the thymus thus unable to produce T cells (69) which is required to fend off robust growth of implanted tumors. Notably, we found that all inoculated MC38 H-2K^d^ tumors grew progressively in immunodeficient BALB/c^nu/nu^ mice (Fig. 8B), eliminating the possibility of a loss of tumorigenicity. These data strongly validated that the MC38 H-2K^d^ tumors are inherently capable of *in vivo* growth. Moreover, spontaneous MC38 H-2K^d^ tumor regression in C57BL/6N PD-1 KO and WT mice was confirmed to be specifically induced by the presence of allogeneic class I MHC.

**Figure 8.**
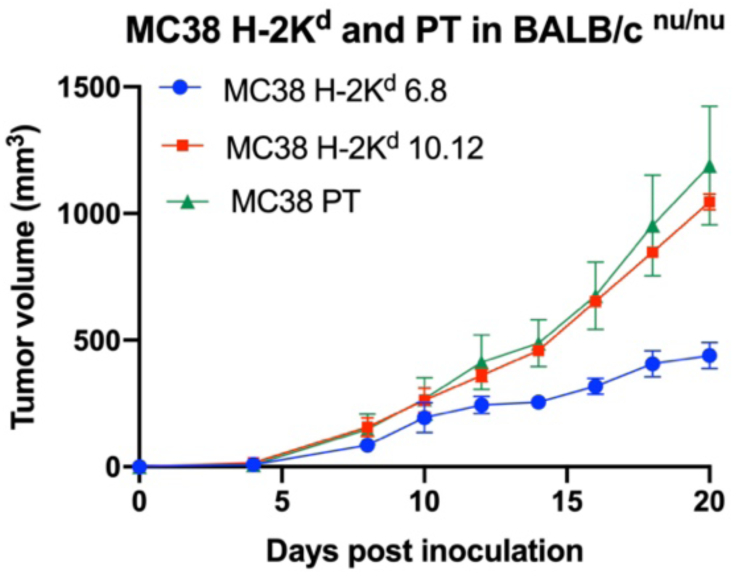
Injection of MC38 tumors into BALB/c^nu/nu^ immunodeficient mice to validate the tumorigenicity of inoculated cells used in this study. Kinetics of tumor growth is shown for MC38 PT, H-2K^d^ clone #6.8, and H-2K^d^ clone #10.12 tumor-bearing hosts. Data represent the means ± SEM of three mice for each group.

### MCA-205 H-2K^d^ tumors regress in WT mice equipped with PD-1

Having established that unmodified MCA-205 cells spontaneously regressed when injected into PD-1 KO animals (Fig. 6A and 6B), we focused on the investigation of MCA-205 H-2K^d^ clones (generated through limiting dilution of initial transfectants) in the WT mice model. Similar to the MC38 study, the MCA-205 H-2K^d^ clone #12.3 was completely regressed (Fig. 9A). Consistently, a different clone of H-2K^d^-overexpressing MCA-205 cancer cells (clone #7.7) also displayed strong antitumor immunity represented by 100% rejection in the WT host (Fig. 9B). These results were confirmed to be achieved through the H-2K^d^-mediated immunity, but not due to the possible immunogenicity of Neo^R^ gene. The MCA-205 Neo^R^ tumors were less visible to the immune cells, resulting in robust growth in the WT animals (Fig. 9C). In addition, the possibility of late tumor progression was excluded in this study as the MCA-205 H-2K^d^-tumor-bearing host did not exhibit any tumor regrowing for about 7 months of observation. Altogether, these data evidently strengthen the hypothesis that allogeneic H-2K^d^ successfully converted the aggressive tumors into highly immunogenic tumors, even in the WT mice equipped with the PD-1/PD-L1 immuno-inhibitory mechanisms.

**Figure 9.**
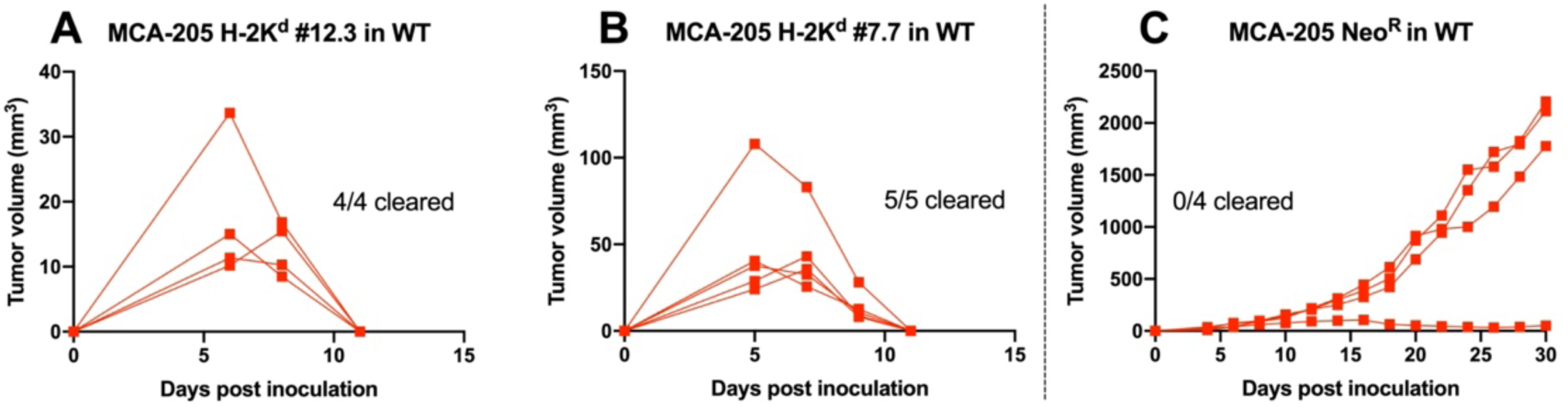
Growth curves of MCA-205 H-2K^d^ tumors indicated spontaneous regression in WT mice. Individual tumor growths of monoclonal MCA-205 H-2K^d^ #12.3 **(A)** and #7.7 **(B)** are shown. **(C)** MCA-205 cells stably transfected with the Neo^R^ gene were used as a control to assess drug resistance-mediated immunogenicity. Data shown are from one experiment, n = 4-5 mice per group.

### Delayed growth of the mixed MC38 H-2K^d^: PT tumors *in vivo*

In the quest of achieving abscopal effect phenomenon, we next asked whether the presence of allogeneic H-2K^d^ cells could affect the immunogenicity against the original MC38 PT cells. To investigate this, prior to subcutaneous inoculation, we mixed an equal cell density (1:1) of MC38 H-2K^d^: PT cells and implanted the mixture into either PD-1 KO or WT mice. The growth of transplanted tumor-mixture was significantly delayed in both mice (Fig. 10A and 10B), contrary to the progressive growth of MC38 PT cell alone. Nevertheless, the speed of *in vivo* growth in WT mice was faster compared to that of PD-1 KO mice. Indeed, the inhibition of immunosuppressive PD-1/PD-L1 signal highly contributed to slow tumor progression in PD-1 KO animals, validating the importance of immune checkpoint blockade. This finding showed that in the pooled mixture of aggressive MC38 PT and immunogenic MC38 H-2K^d^ cells, trans-acting cross-immunity might have occurred against the poorly immunogenic parental cells, which is evidently mediated by the allogeneic H-2K^d^.

**Figure 10.**
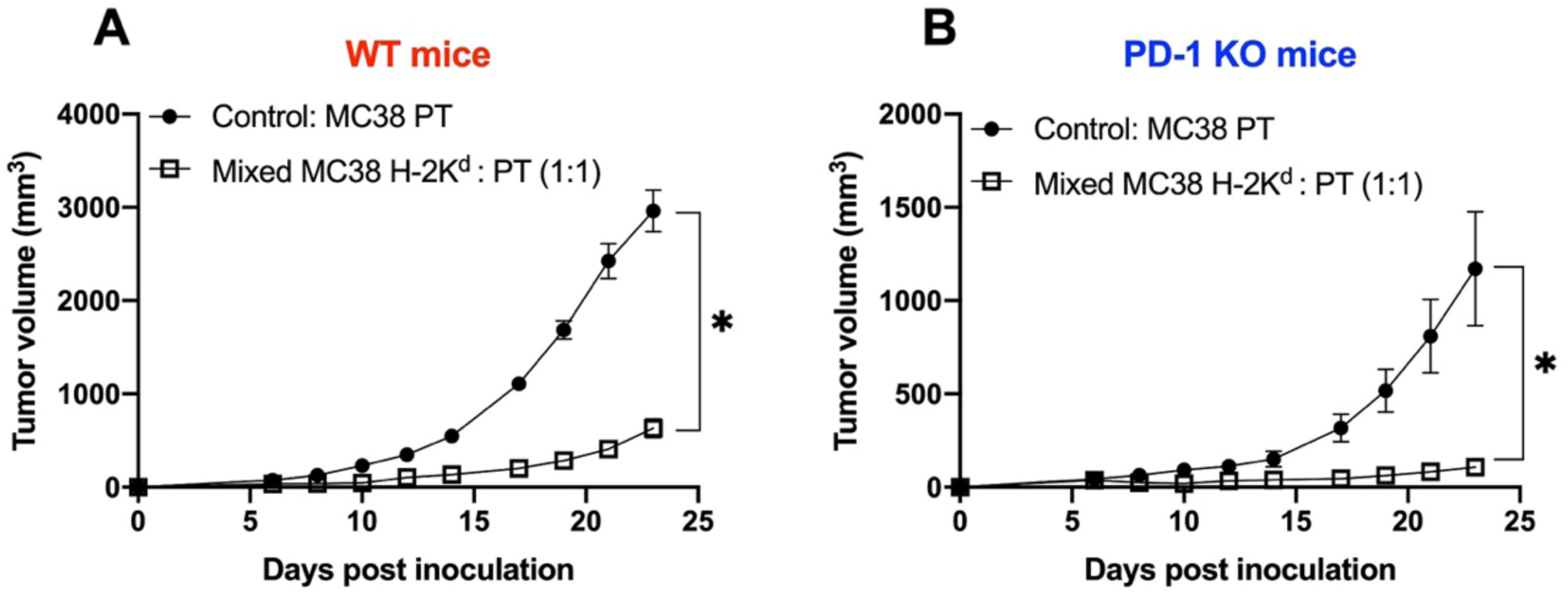
Delayed growth of the mixed MC38 H-2K^d^: PT tumors *in vivo*. Prior to injection into PD-1 KO and WT mice, the equal numbers of MC38 PT cells (5×10^5^ cells/100 µl) and MC38 H-2K^d^ clone #6.8 (5×10^5^ cells/100 µl) were counted and mixed, totaling the final cell concentration to 5×10^5^ cells/100 µl per inoculation. As a control, MC38 PT cells (5×10^5^ cells/100 µl) were injected into both mice. Palpable tumor growth was then measured once in every 2 days. The mixed-tumor growth in WT mice **(A)** and in PD-1 KO mice **(B)** showed delayed progression compared with MC38 PT tumor inoculated with similar total cell density. Data shown are from one experiment, n = 4-5 mice per group. Average tumor growth curves depict means ± SEM. Some error bars are shorter than the size of the symbol hence cannot be shown. Statistical significance was determined by unpaired Student’s *t*-test.

### Sensitizing PD-1 KO mice with immunogenic H-2K^d^ tumors results in protection against parental MC38 challenge

Next, we sought to determine the extent to which H-2K^d^-overexpressing tumors could protect the mice against unmodified cancers. The MC38 H-2K^d^ clones were further investigated in the settings of concomitant tumor immunity and the possibilities of eliciting the abscopal effect. Concomitant tumor immunity is a phenomenon in which a tumor-bearing host is resistant to the growth of similar secondary tumor implants at the distant site (70,71). To test this, we developed sensitization strategies by inoculating MC38 H-2K^d^ tumors (clone #6.8) into the flank of PD-1 KO mice followed by challenging them with parental cells on the contralateral side after 6 days (Fig. 11A). Surprisingly, the sensitization treatment provided complete protection against the unmodified MC38 tumor challenge with all mice rejecting the tumor inoculums (Fig. 11B). Reproducibly, inoculating a different clone of MC38 H-2K^d^ tumors (clone #10.12) resulted in parental tumor rejection in all tested PD-1 KO mice (Fig. 11C).

**Figure 11.**
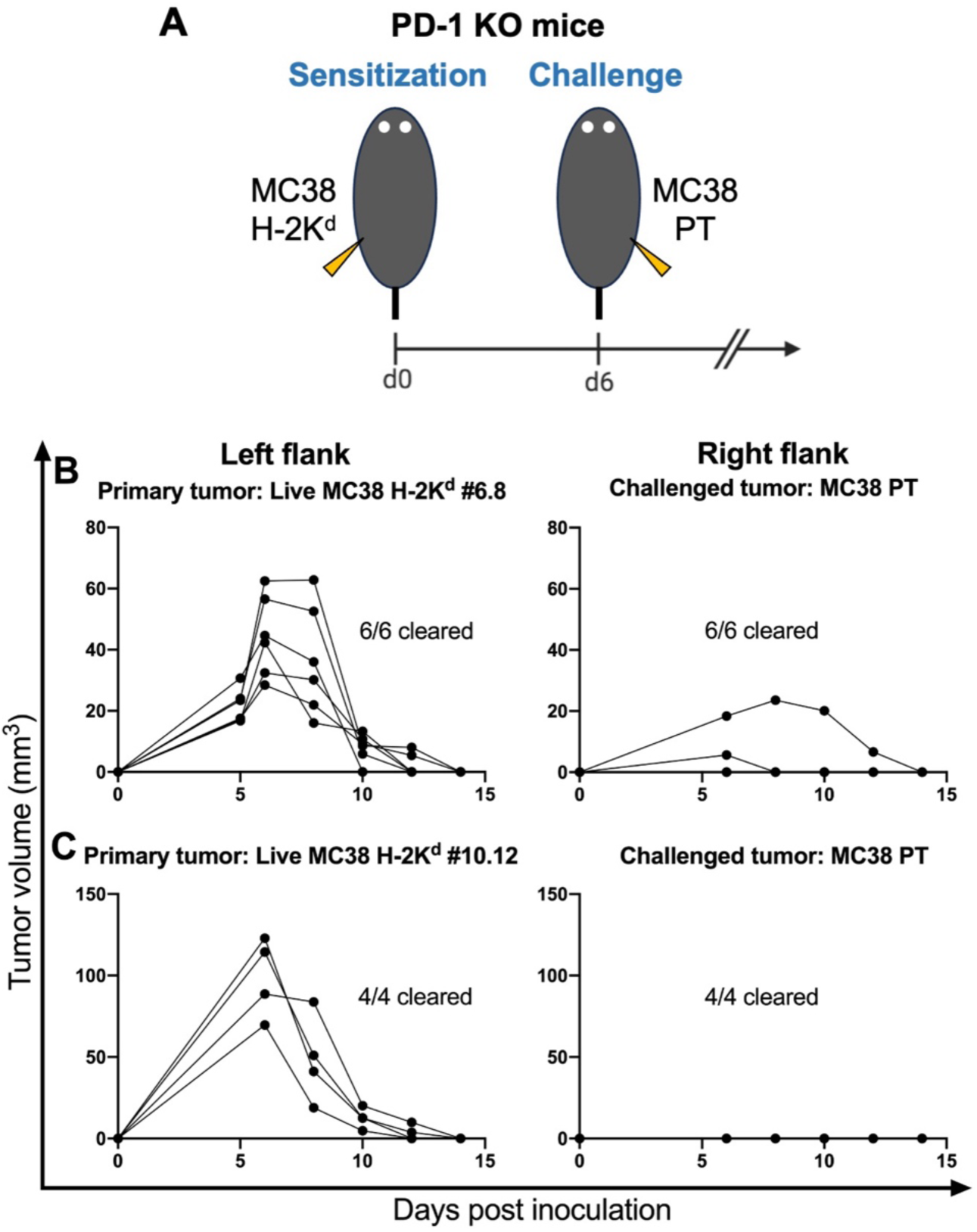
MC38 H-2K^d^ induces protective immunity against parental cell challenge. **(A)** Experimental design: PD-1 KO mice were implanted with MC38 H-2K^d^ on the left flank followed by challenge on the contralateral side with parental MC38 PT on day 6. **(B)** Individual growth curves of primary and secondary tumors of MC38 H-2K^d^ #6.8-sensitized mice. **(C)** Another clone of MC38 H-2K^d^ (clone #10.12) was tested in a similar fashion. Data are representative of two independent study repeats with n = 4-6 mice per group.

Remarkably, sensitizing PD-1 KO mice with growth-arrested (MMC-treated) cells or debris of both clones of MC38 H-2K^d^ tumors provided full protection against the encounter of secondary unmodified tumors (Fig. 12A-C). This finding implied that MC38 H-2K^d^ retained high immunogenicity regardless of their compromised growing capacity and dead/alive status. Most interestingly, sensitized animals immediately gained immunity against the challenge of secondary parental tumors at the distant site even without employing any additional immune adjuvants. This implies that H-2K^d^ itself could act as an adjuvant to boost antitumor immunity.

**Figure 12.**
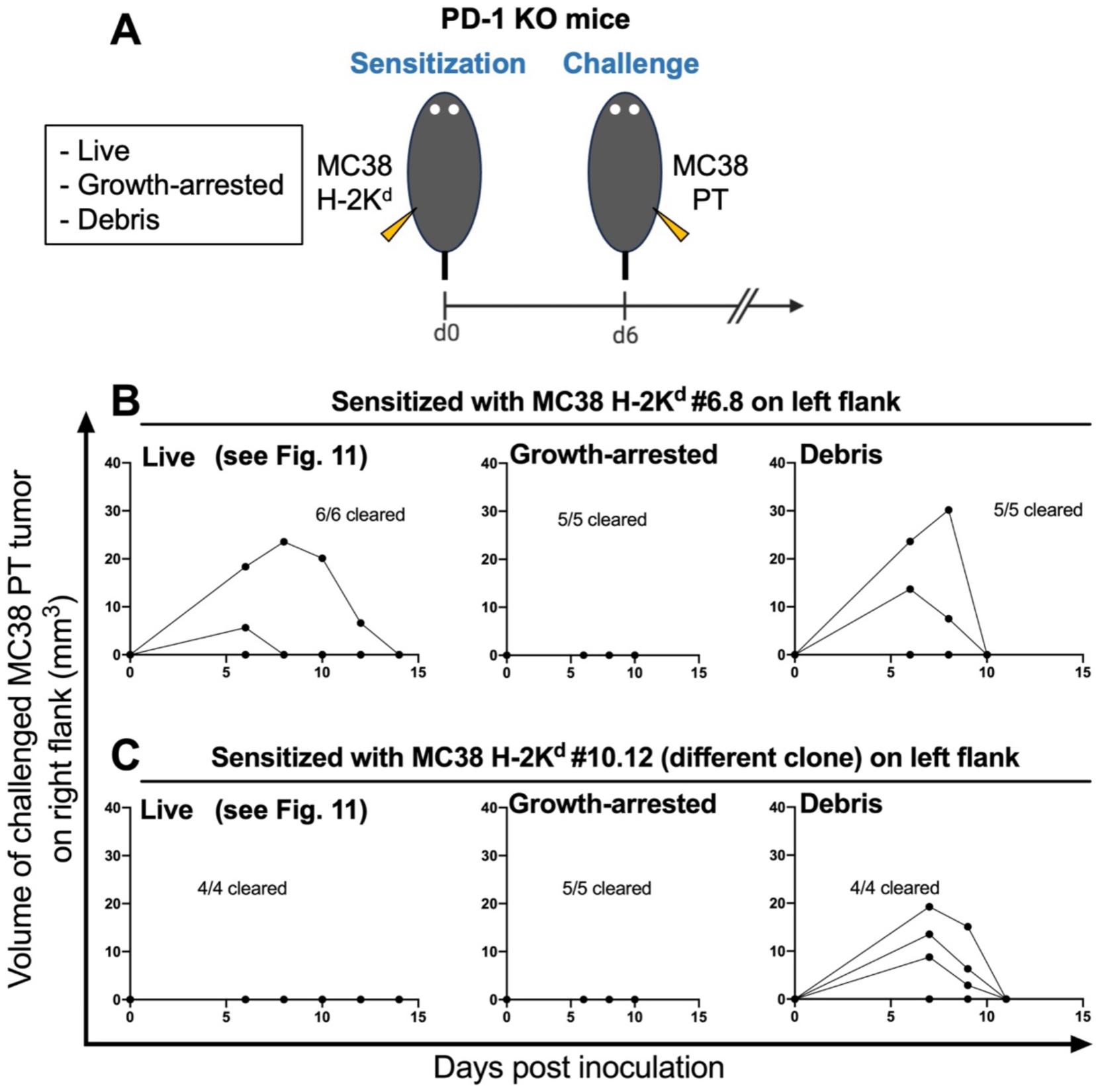
Kinetics of the secondary tumor growth (MC38 PT) in PD-1 KO mice sensitized with MC38 H-2K^d^. **(A)** Experimental design: three different conditions of immunogenic MC38 H-2K^d^ tumors; live, growth-arrested (Mitomycin C-treated), and debris (frozen-thawed cells) were inoculated in the sensitization experiment settings, and on day 6, MC38 PT cells were injected subcutaneously on the contralateral side. Individual tumor growth curves of the challenged MC38 PT tumors in PD-1 KO sensitized with live, growth-arrested, and debris of MC38 H-2K^d^ clones #6.8 **(B)** and #10.12 **(C)**. Data shown are from two experiment with n = 4-6 mice per group.

### MC38 PT cell debris fails to induce protective immunity

Having established an effective sensitization treatment with MC38 H-2K^d^ in PD-1 KO mice, we further asked whether this phenomenon was specifically due to the H-2K^d^– mediated immune modulation rather than the general effects of the MC38 cell itself. To clarify this point, we pre-sensitized PD-1 KO mice with MC38 PT cells (live, growth-arrested, debris) followed by observation of challenged MC38 PT tumor growth in the distant site (MC38 PT vs. MC38 PT, as shown in Fig. 13A).

**Figure 13.**
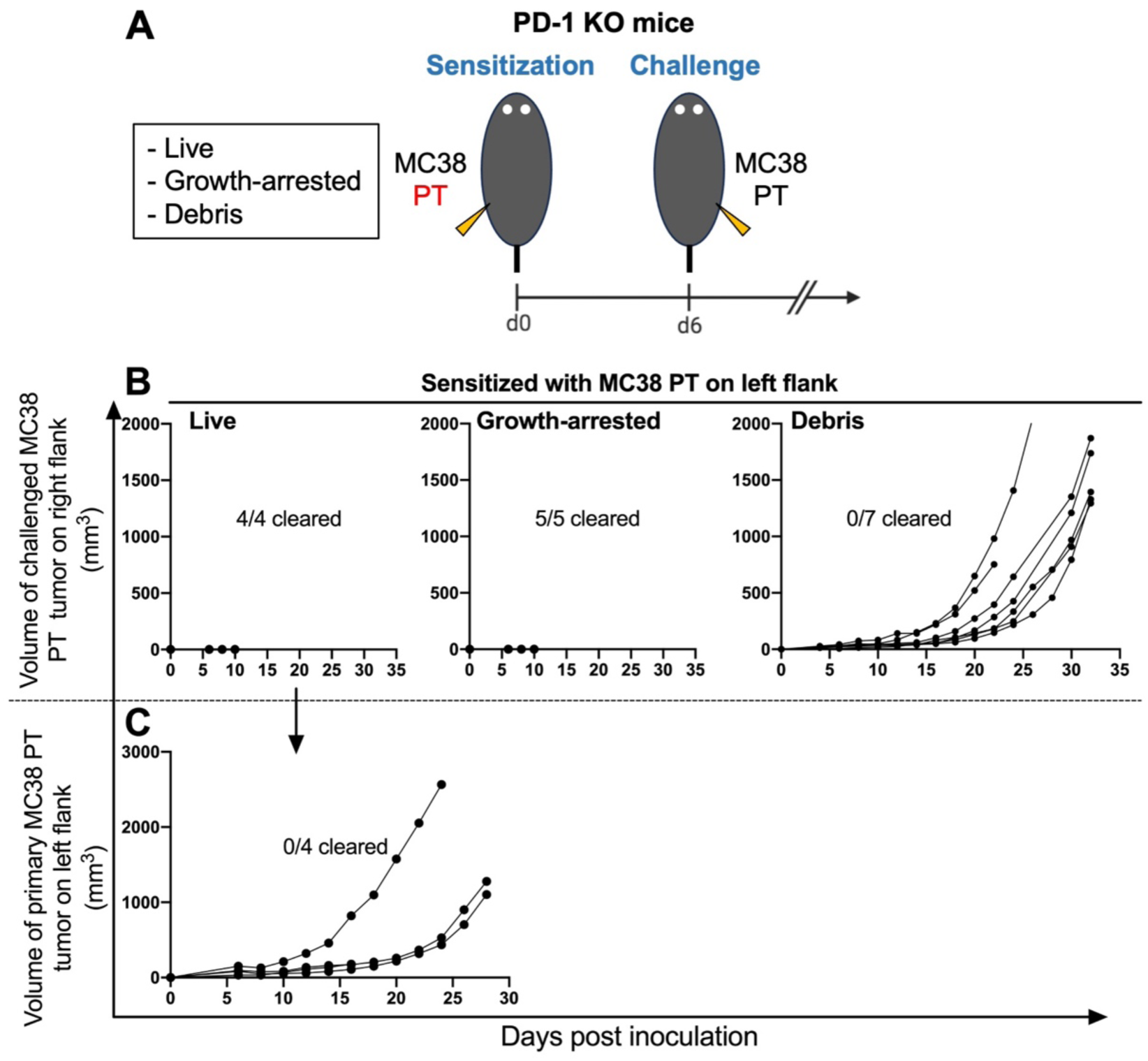
**(A)** MC38 PT tumors were treated (live, growth-arrested, and debris) and inoculated into PD-1 KO mice as a sensitization regimen. Mice were challenged six days later with the same MC38 PT cell line. **(B)** Summaries of the challenged MC38 PT tumor growth. Representative data are shown as individual tumor growth curves. Data for sensitization with MC38 PT cell debris are pooled from two independent experiments. **(C)** Live MC38 PT cells were implanted as the primary tumor in order to sensitize PD-1 KO mice. Data shown as individual tumor growth curves from one independent experiment with n = 4.

Sensitization (of PD-1 KO mice) with growth-arrested MC38 PT cells provoked antitumor immunity wherein all mice rejected inoculums in both flanks effectively (Fig. 13B). Moreover, concomitant immunity occurred when PD-1 KO mice received primary inoculation of live MC38 PT, indicated by a complete rejection of equivalent live MC38 PT tumor (Fig. 13B). However, due to the robust growth of primary tumor, the subjected mouse condition was deteriorated rapidly and ultimately resulted in a faster endpoint of the experiment (Fig. 13C), in comparison with the long-term tumor-free status of animals sensitized with the highly immunogenic H-2K^d^ tumor (Fig. 12).

Most excitingly, we found that sensitization with cell debris of MC38 PT failed to prevent tumor growth, evidently displayed by the progressive growth of challenged live parental cells (Fig. 13B). These findings further highlight the ability of MC38 H-2K^d^ debris, but not MC38 PT debris, to mount a markedly effective immunity, particularly in the absence of PD-1 activity.

### Sensitization with MC38 H-2K^d^ cell debris provides long-term immunological memory against MC38 carcinoma

When taking into account that sensitizing PD-1 KO mice with MC38 H-2K^d^ prompted a strong anti-tumor immunity, we next sought to validate whether this treatment could be sufficient to protect sensitized mice from subsequent tumor rechallenge in a long period of time. To more deeply assess long-term protective immunity, about 5 months following the sensitization, all mice that had survived the first tumor (MC38 PT) inoculation were rechallenged with MC38 PT on the same site as the first injection (Fig. 14A). Observation of palpable tumor formation was conducted for a period lasting more than 8 weeks to exclude the possibility of delayed onset of tumor growth.

**Figure 14.**
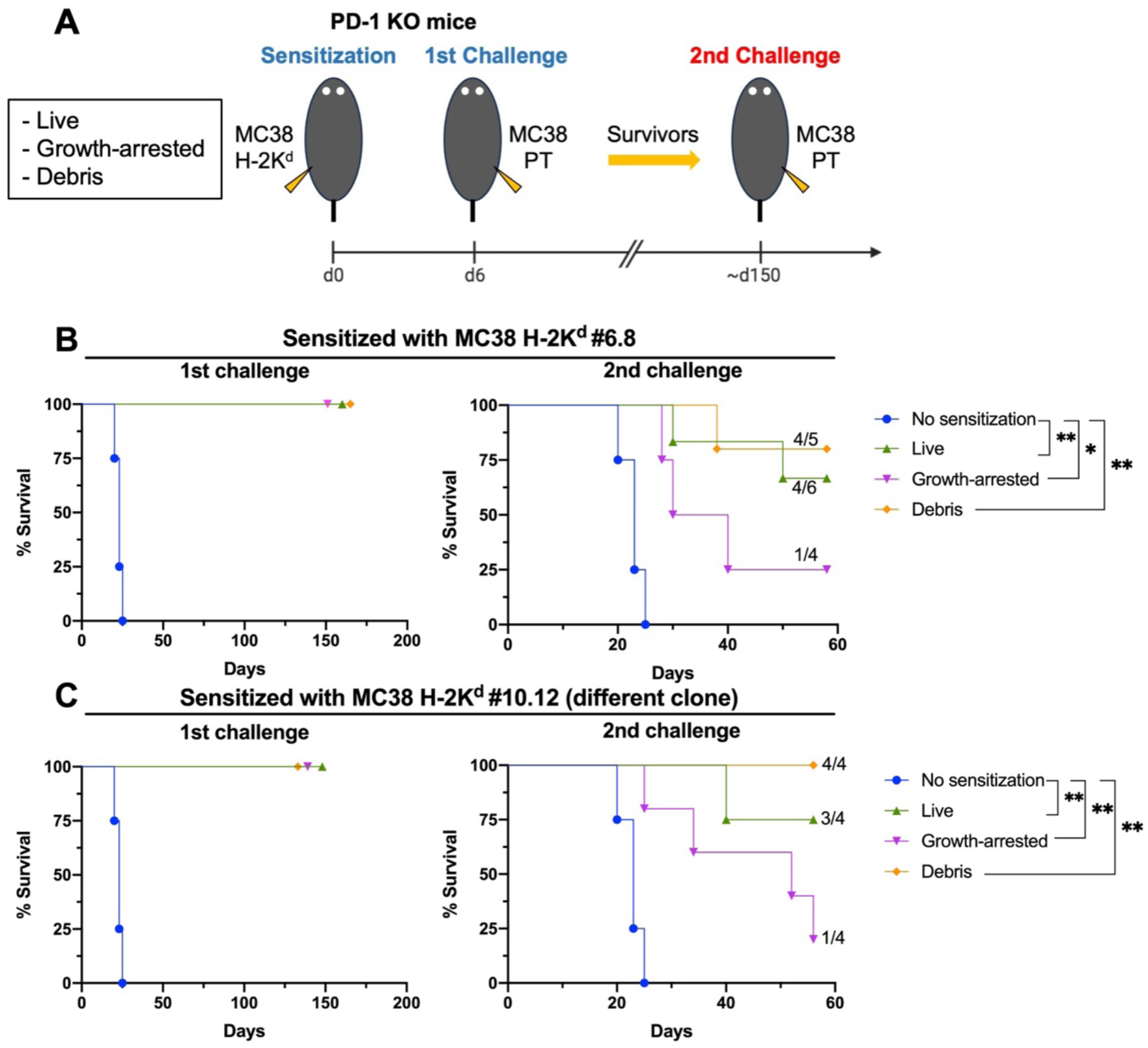
**(A)** Experimental design: PD-1 KO mice were sensitized on the left flank with each clone of MC38 H-2K^d^ cells (live, growth-arrested, debris) and on day 6, MC38 PT were injected subcutaneously on the contralateral side. Upon the first challenge, tumor growth was recorded and mice that rejected tumors were rechallenged around 150 days later with MC38 PT. (**B** and **C**) Kaplan-Meier survival curves following the first and second tumor challenges in PD-1 KO mice pre-sensitized with MC38 H-2K^d^ #6.8 (B) and MC38 H-2K^d^ #10.12 tumors (C). Naïve PD-1 KO mice were used as positive controls for tumor growth (no sensitization). Endpoint defined as tumor volume > 1,000 mm^3^. Data shown are from one experiment, n = 4 to 6 mice per group. Statistical significance was determined by log-rank (Mantel-Cox) test.

Interestingly, PD-1 KO mice that were sensitized with the debris of MC38 H-2K^d^ cells showed the strongest immune response, with 80 % (MC38 H-2K^d^ #6.8–sensitized) and 100 % (MC38 H-2K^d^ #10.12–sensitized) of the mice surviving the second tumor challenge (Fig. 30B and 30C). In contrast, rejection rates of growth-arrested–sensitized mice from both H-2K^d^ clones reproducibly showed the lowest survival, indicating failure to induce long-term immunity (Fig. 15). Furthermore, treatment with live MC38 H-2K^d^ cells also showed durable memory response with more than 60% of rechallenged mice eliminated the tumors, however, in the practical settings, administrating live tumor cells as prophylaxis is ethically impossible. Taken together, these data put an emphasis that the greater and durable immune responses were significantly induced by MC38 H-2K^d^ cell debris, particularly in the absence of PD-1 activity.

**Figure 15.**
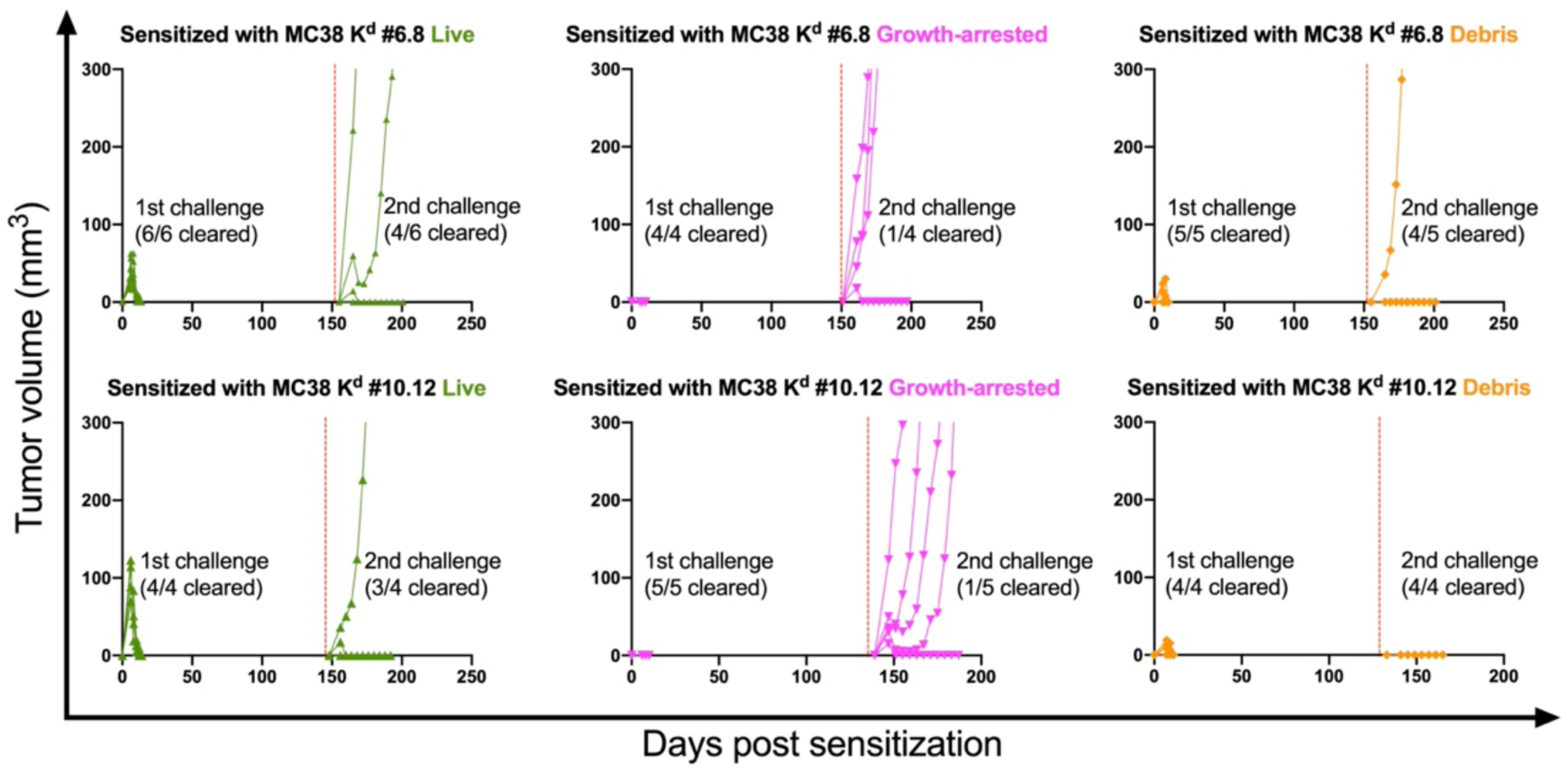
PD-1 KO mice were implanted with live (green), growth-arrested (magenta), and cell debris (orange) of MC38 H-2K^d^ clone #6.8 or clone #10.12 tumor followed by the first challenge with MC38 PT. All survivors were rechallenged (red dashed line) 5 months later with MC38 PT cell line at the same site as the first challenge. Representative data are shown as individual tumor growth curves.

### Absence of PD-1 is required for successful rejection of challenged MC38 PT

Having elucidated the potent induction of anti-tumor immunity in PD-1 KO mice, we asked whether MC38 H-2K^d^ cells could also sensitize WT mice for the protection against challenged MC38 PT tumor *in vivo.* To formally address this question, the sensitization regimen was then tested in WT mice in which the PD-1/PD-L1 axis functions normally (Fig. 16A). In the presence of the PD-1 inhibitory receptor, antitumor immunity against parental MC38 cells was diminished as indicated by the robust tumor growth (Fig. 16B, right flank). Furthermore, the unanticipated finding was unveiled by the regrowing of MC38 H-2K^d^ tumor in WT mice (Fig. 16B, left flank), necessitating the importance of PD-1 blockade to reinvigorate host immunity against tumor. Clearly, by revisiting the successful sensitization results in PD-KO mice wherein both H-2K^d^ and PT tumors were completely rejected (Fig. 16C), these findings strongly identify the synergistic effect of PD-1 blockade and allogenic H-2K^d^ in the tumor niche.

**Figure 16.**
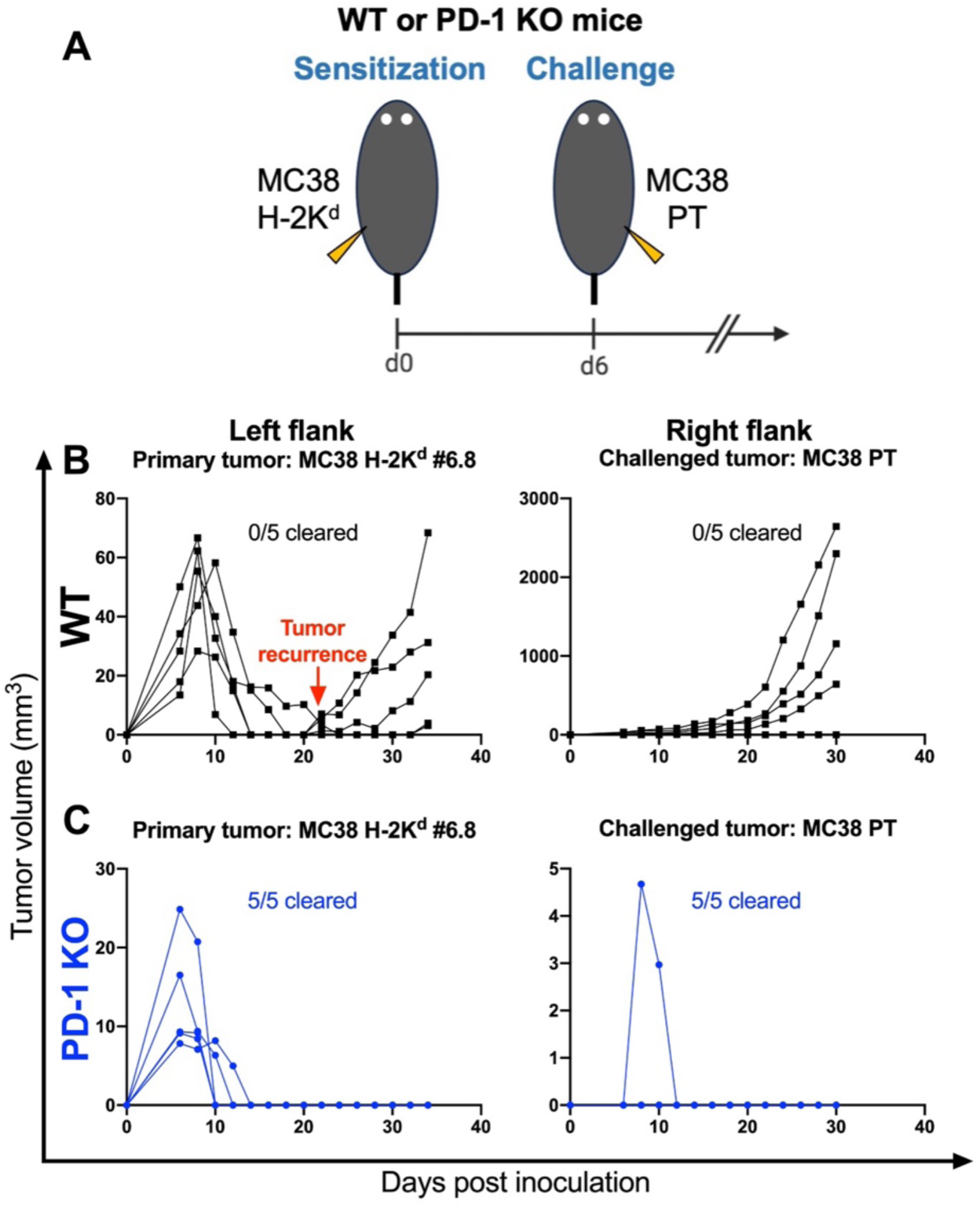
**(A)** Experimental design: WT or PD-1 KO mice were sensitized with live MC38 H-2K^d^ and then challenged with MC38 PT on day 6. **(B)** Kinetics of primary tumor growth (MC38 H-2K^d^) in WT mice (black line) showed initial tumor rejection followed by recurrence, while on the opposite flank, the challenged parental line grew robustly. **(C)** Growth curves of individual tumors showing complete rejection of both tumors in PD-1 KO mice (blue line) as a comparison. Data shown are from two repeated experiments, n = 4-6 mice per group.

Next, to provide a proportional comparison with sensitization study in PD-1 KO mice, WT animals were also injected with inactivated or dead tumor cells ectopically expressing H-2K^d^ 6 days prior to parental tumor challenge (Fig. 17A). The secondary MC38 PT tumors were partially rejected in WT mice pre-sensitized with live (2/4 mice) and growth-arrested (1/4 mice) MC38 H-2K^d^ cells (Fig. 17B). Moreover, WT mice that received debris of MC38 H-2K^d^ cells as primary inoculums completely failed to acquire sensitivity to the challenged parental MC38 tumor rejection in comparison with that of PD-1 KO mice. These findings corroborated the fact that the absence of PD-1 inhibitory signal is a prerequisite to generate protective antitumor immunity against the aggressive parental cell line.

**Figure 17.**
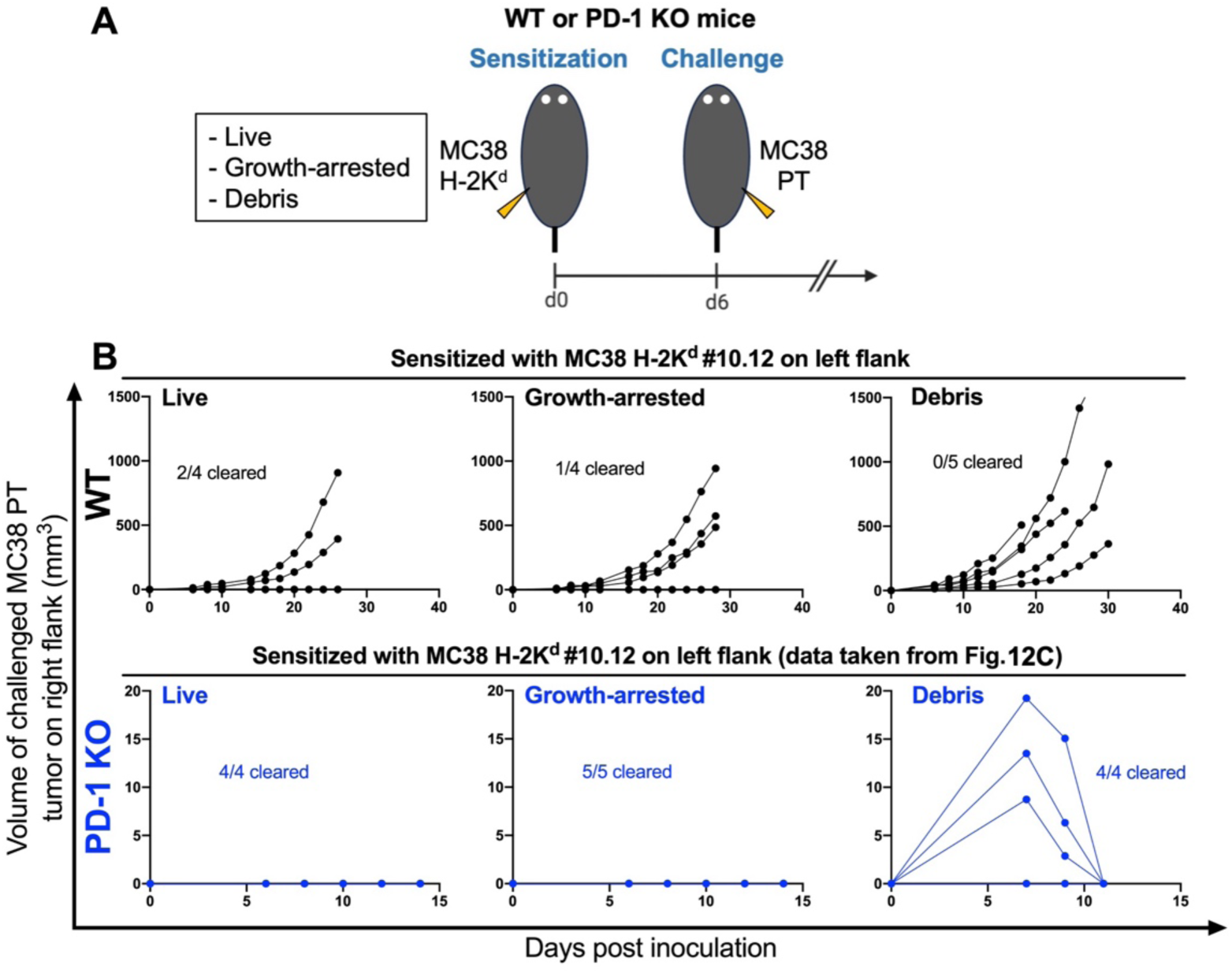
**(A)** Experimental design: WT or PD-1 KO mice were sensitized with MC38 H-2K^d^ (live, growth-arrested, and debris) and on day 6, MC38 PT was injected subcutaneously on the contralateral side. **(B)** Individual tumor growth kinetics of parental MC38 tumor in WT animals (black) versus PD-1 KO mice (blue). Complete tumor rejection was achieved in all PD-1 KO mice while WT mice failed to completely eradicate the challenged tumor.

### Increased PD-1 expression in CD8^+^ T cells alleviates anti-tumor immune responses

To validate the importance of the absense of the PD-1 axis to sustain anti-tumor immunity, the tumor-infiltrating lymphocytes (TILs) were isolated from the MC38 H-2K^d^–sensitized WT mice bearing the secondary MC38 PT tumor. Indeed, a marked increase in the frequencies of CD8^+^ PD-1^+^ T cells was detected (Fig. 18A). However, there was no significant difference in the percentage of CD8^+^ PD-1^+^ T cells between sensitization treatments (Fig. 18B). High PD-1 expression within the infiltrating CD8^+^ T cells is deemed to poor prognosis (72, 73) due to the inactivation of effective tumor-killing ability. Overall, these data strengthen the importance of the immunomodulation effects of PD-1 deficiency in synergizing with H-2K^d^–induced antitumor immunity.

**Figure 18.**
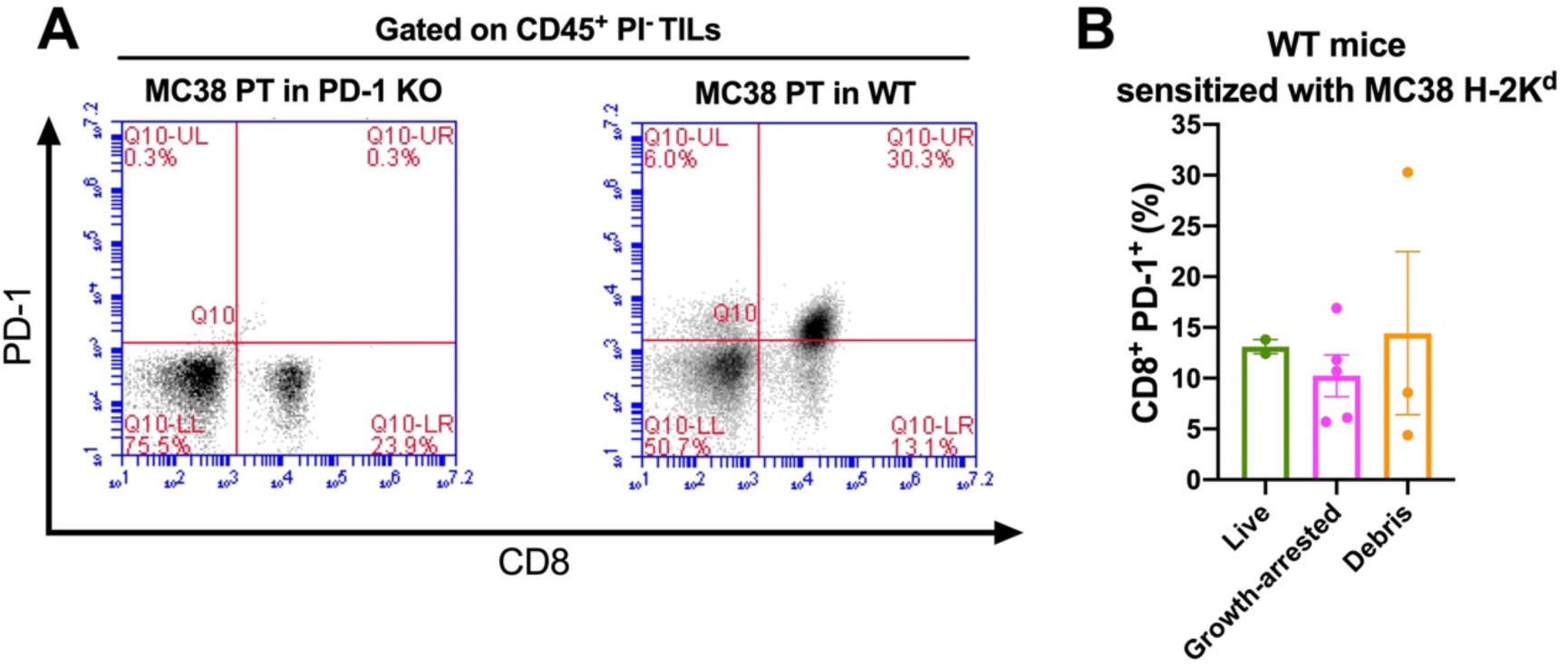
Elevated PD-1 expression in CD8^+^ T cells from tumor-bearing WT mice. **(A)** Representative FACS patterns of CD8^+^ PD-1^+^ cells in tumors isolated from PD-1 KO (control) and WT mice. TILs after tumor digestion were stained with anti-CD45 and propidium iodide (PI) in order to gate live lymphocytes. The established gate was then plotted to identify CD8 and PD-1 cells population. **(B)** Bar graphs of CD8^+^ PD-1^+^ frequency among TILs of MC38 H-2K^d^–sensitized WT mice. No statistical significance was achieved.

### The specific immunity against tumor antigens is required to elicit antitumor responses

As shown above, the H-2K^d^-harboring tumor has completely protected the animals from the challenge of unmodified tumors. Next, we asked whether the retardation of tumor growth is also specific to tumor antigens rather than a universal effect of allogeneic H-2K^d^ presentation. To assess this question, we established a concomitant immunity model between MCA-205 and MC38 cell lines. In this fashion, the mice received administration of secondary tumor inoculation that was distinct from the preexistent primary tumor (*e.g.,* MC38 as primary tumor and MCA-205 as secondary tumor or vice versa).

In PD-1 KO, it was already shown that the parental MC38 grew robustly but not when the mice were firstly sensitized by MC38 overexpressing H-2K^d^ (Fig. 11). In contrast, the PD-1 KO that had received MCA-205 H-2K^d^ cells as a primary tumor was not protected against subsequent MC38 tumor challenge in the distant site (Fig. 19A), indicating that the triggered immune reaction was due to MC38-specific tumor immunity.

**Figure 19.**
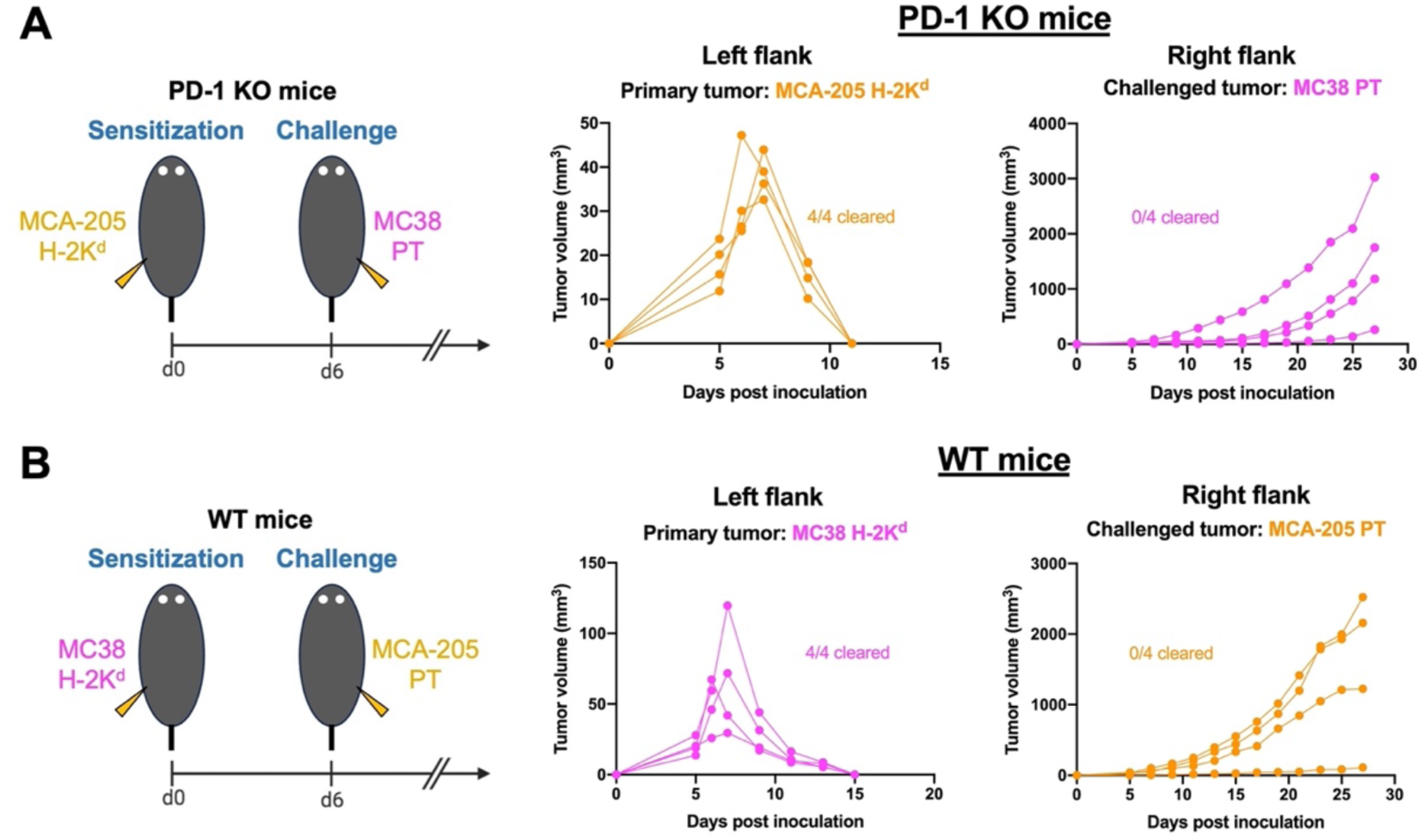
Individual tumor growth curves of criss-cross experiments between MC38 and MCA-205 cancer cells. **(A)** PD-1 KO mice were subcutaneously inoculated with MCA-205 H-2K^d^ as the primary tumor and then challenged with unmodified MC38 cells at day 6. **(B)** WT mice were injected with inoculums of MC38 H-2K^d^ and received a different parental line (MCA-205) as the secondary tumor. Data shown are from one experiment, n = 4 mice per group.

A similar experiment using WT mice to test the MCA-205-specific immunity was conducted. Because MCA-205 readily formed tumors in WT mice (Fig. 19), we inoculated these mice with MC38 H-2K^d^ tumors and sequentially challenged them with the MCA-205 parental line. As anticipated, although the immunogenic MC38 H-2K^d^ tumors spontaneously regressed, the inoculated mice were naïve for MCA-205-derived tumor antigens hence they could not exhibit immunity against the sarcomas (Fig. 19B). Having observed the phenomena of specific immunity, it is evident that the display of tumor-specific antigens through MHC class I are essential for successful antitumor immunity.

### Infiltration of CD11c^+^ and CD8^+^ cells is observed in the injection sites of MC38 H-2K^d^

Evaluation of infiltrating immune cells in MC38 tumors ectopically expressing H-2K^d^ was impractical to conduct because the tumors reached their maximum size between day 6 and 10 and spontaneously regressed thereafter (Fig. 7A and 7B). In addition, the size of palpable tumors was very small, adding constraints to isolate bulk tumors for downstream analysis. Similarly, it was impossible to locate the inoculated MC38 H-2K^d^ cell debris *in vivo*. Since sensitization with the debris of MC38 H-2K^d^ generated durable antitumor immune responses, analysis of infiltrating immune cells becomes exciting to accomplish.

To more deeply investigate the possible involvement of immune cells inside the highly immunogenic tumor microenvironment, we embedded MC38 H-2K^d^ tumors or their cell debris in the Matrigel basement membrane. Matrigel solidifies at body temperature, resulting in a plug that might trap the infiltrating host cells. In this study, Matrigel plugs containing MC38 H-2K^d^ cells (live or debris) were isolated from PD-1 KO mice on day 7 and were visualized by hematoxylin and eosin (HE) staining and immunofluorescence staining.

HE staining of Matrigel carrying live MC38 H-2K^d^ cells showed the massive number of nucleated cells within the plug (Fig. 20A and 20B). Possible penetration of host immune cells toward the Matrigel plug was suggested by the presence of nucleated cells nearby blood vessels at the periphery side of the plug (Fig. 20B). Furthermore, precise localization of cell debris *in vivo* after injection was made possible by embedding dead MC38 H-2K^d^ cells in Matrigel (Fig. 20C and 20D). Histological analysis revealed remarkably fewer nucleated cells in the MC38 H-2K^d^ debris-containing Matrigel plug (Fig. 20C). At day 7 after inoculation, some nucleated cells were seen trapped within the Matrigel, near to the subcutaneous tissue (Fig. 20D). This solid disparity of histological visuals between live and dead MC38 H-2K^d^ cells inside the Matrigel plug has validated the de facto situation of tumor microenvironment.

**Figure 20.**
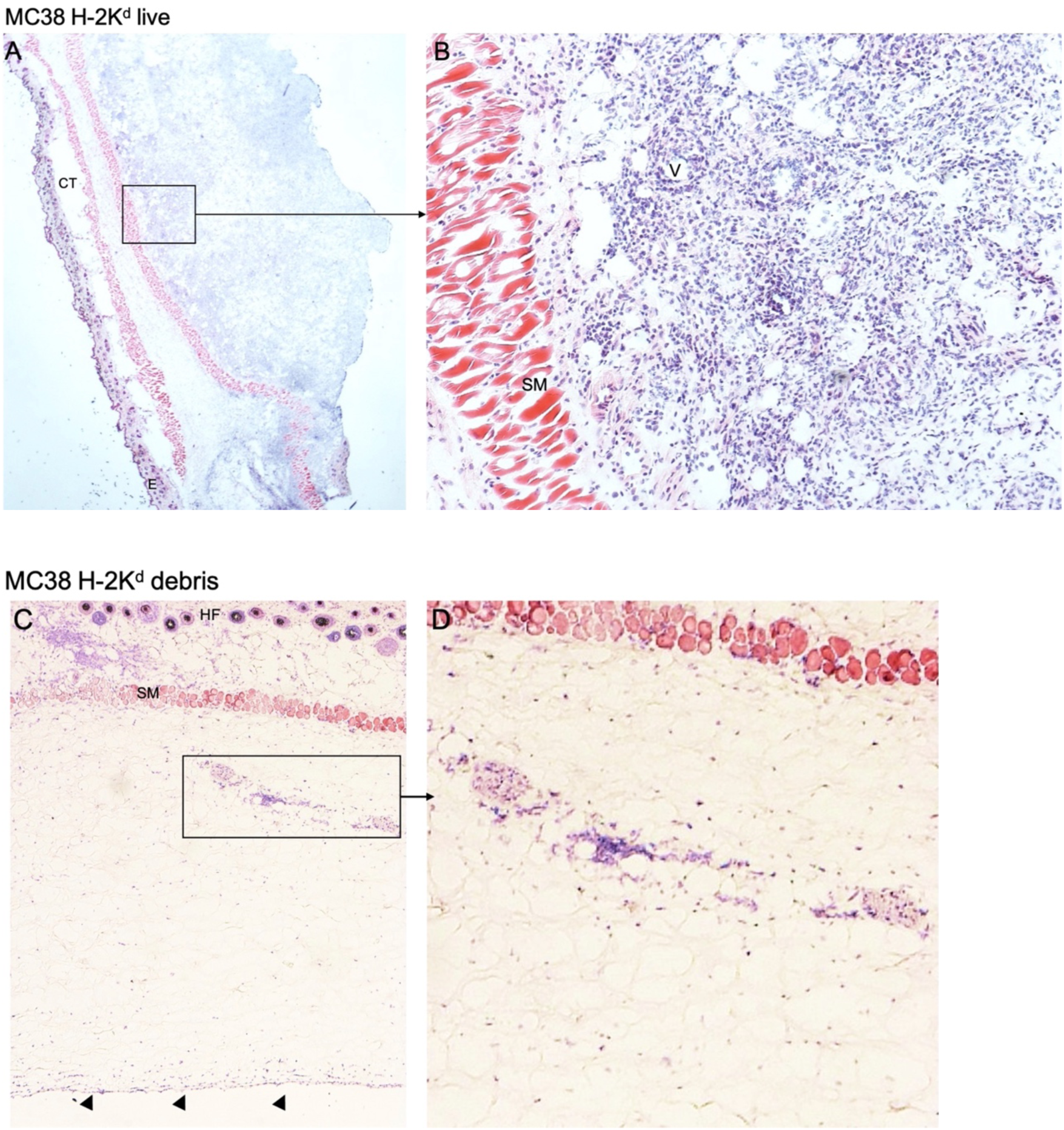
Representative histological images of HE-stained Matrigel plugs containing live MC38 H-2K^d^ cells **(A** and **B)** showing the large number of nucleated cells within the Matrigel area at 7 days after injection. (B) Higher magnification of the area in (A) shows the blood vessel and possible penetration of host cells. **(C** and **D)** Subcutaneous Matrigel plug containing MC38 H-2K^d^ cell debris showing the border of Matrigel plug indicated by black arrowheads (C) and a remarkably smaller number of nucleated cells within Matrigel plug (D) in comparison with that of live cells (B). CT, soft connective tissue; SM, striated muscle; E, epidermis; HF, hair follicle; V, blood vessel.

Next, in order to distinguish between tumor cell population and infiltrating host cells, the live MC38 H-2K^d^ cells-containing Matrigel sections were further immunostained. The nucleated cells (DAPI-stained) inside the Matrigel plug can be divided into MC38 H-2K^d^ tumor cells and penetrating host immune cells that were stained well with markers for cytotoxic T lymphocytes (CD8α) and dendritic cells (CD11c) (Fig. 21A). Large numbers of CD11c^+^ and CD8^+^ host cells were seen infiltrating the Matrigel plug, of which several cells were in close contact, pinpointing the possibility of cross-presentation (Fig. 21B and 21C). Professional antigen-presenting cells (APC) such as dendritic cells have the ability to capture and to engulf tumor cells and present tumor antigens to CD8^+^ T cells, a process referred to as cross-presentation (37).

**Figure 21.**
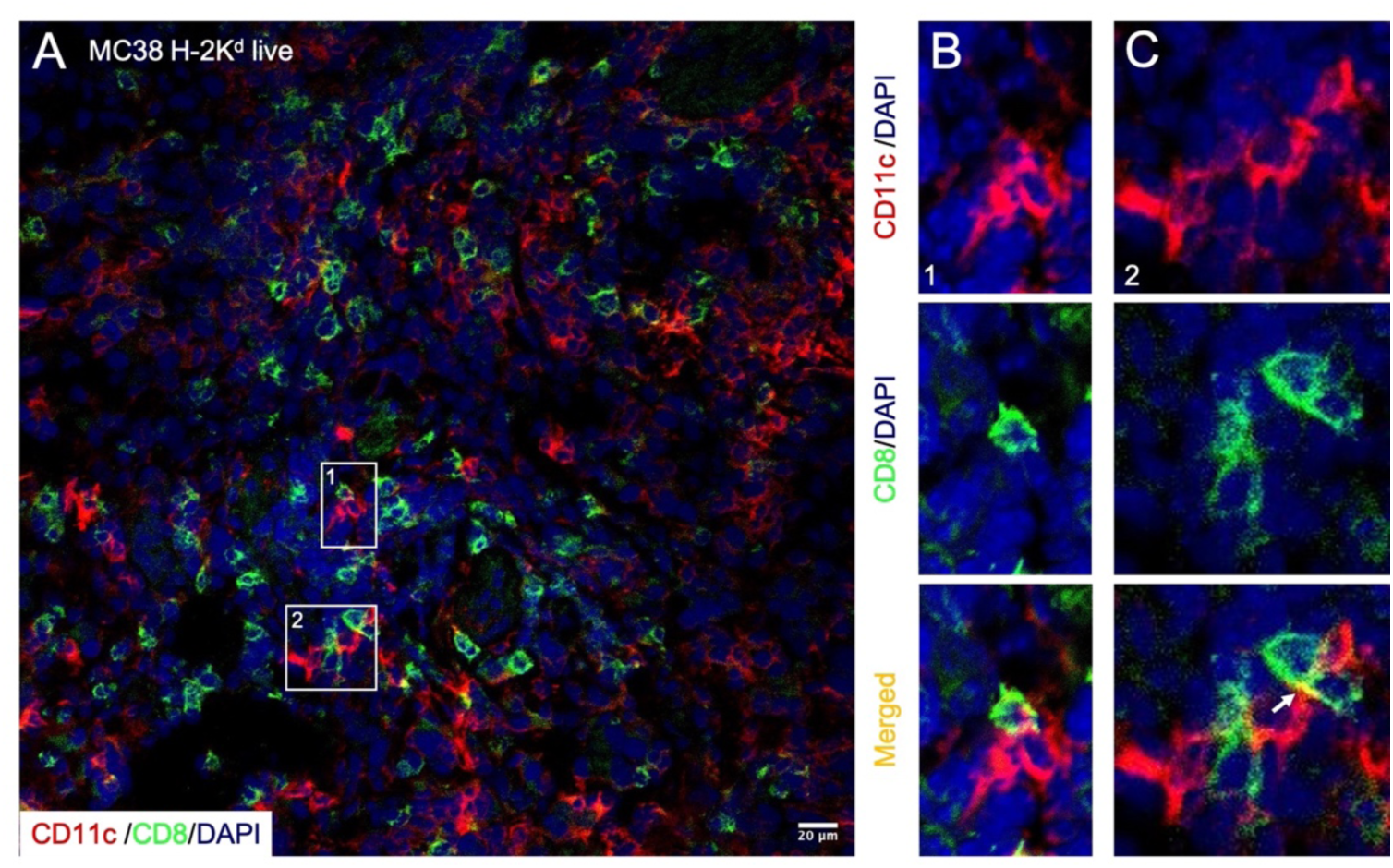
Infiltration of host immune cells into Matrigel plugs containing MC38 H-2K^d^ cells. PD-1 KO mice were injected subcutaneously in the flank with live MC38 H-2K^d^ cells embedded in Matrigel and plugs were isolated for immunohistochemistry 7 days after injection. **(A** to **C)** Representative images of plug sections immunostained for CD11c^+^ (dendritic cells, red) and CD8^+^ (T cells, green), and 4′,6-diamidino-2-phenylindole (DAPI) (nuclei, blue). **(A)** Host cells expressing CD11c and CD8 are present within the Matrigel area. **(B** and **C)** Higher magnification of the area inside the region of interest marked with the white box in (A). White arrow shows possible contact between CD11c^+^ cells and CD8^+^ cells.

Last, we observed massive infiltration of CD11c^+^ cells in Matrigel plugs harboring debris of MC38 H-2K^d^ cells (Fig. 22A). Consistently, direct cell-cell contact between CD11c^+^ and CD8^+^ cells was also observed in Matrigel plugs containing the debris of MC38 H-2K^d^ cells (Fig. 22B and 22C). Apparently, the efficient uptake of tumor antigens by APC might be associated with positive anti-tumor immune responses in the sensitization (Fig. 12B-C) and the tumor rechallenge experiments (Fig. 14B-C) in this study. Therefore, the mingling of CD11c–CD8 in the tumor microenvironment might serve as an indicator for a favorable prognosis. Notably, the recruitment of host immune cells was dependent on the presence of cancer cells and not caused by the Matrigel itself since very few cells were seen infiltrating cell-free Matrigel plug (data not shown).

**Figure 22.**
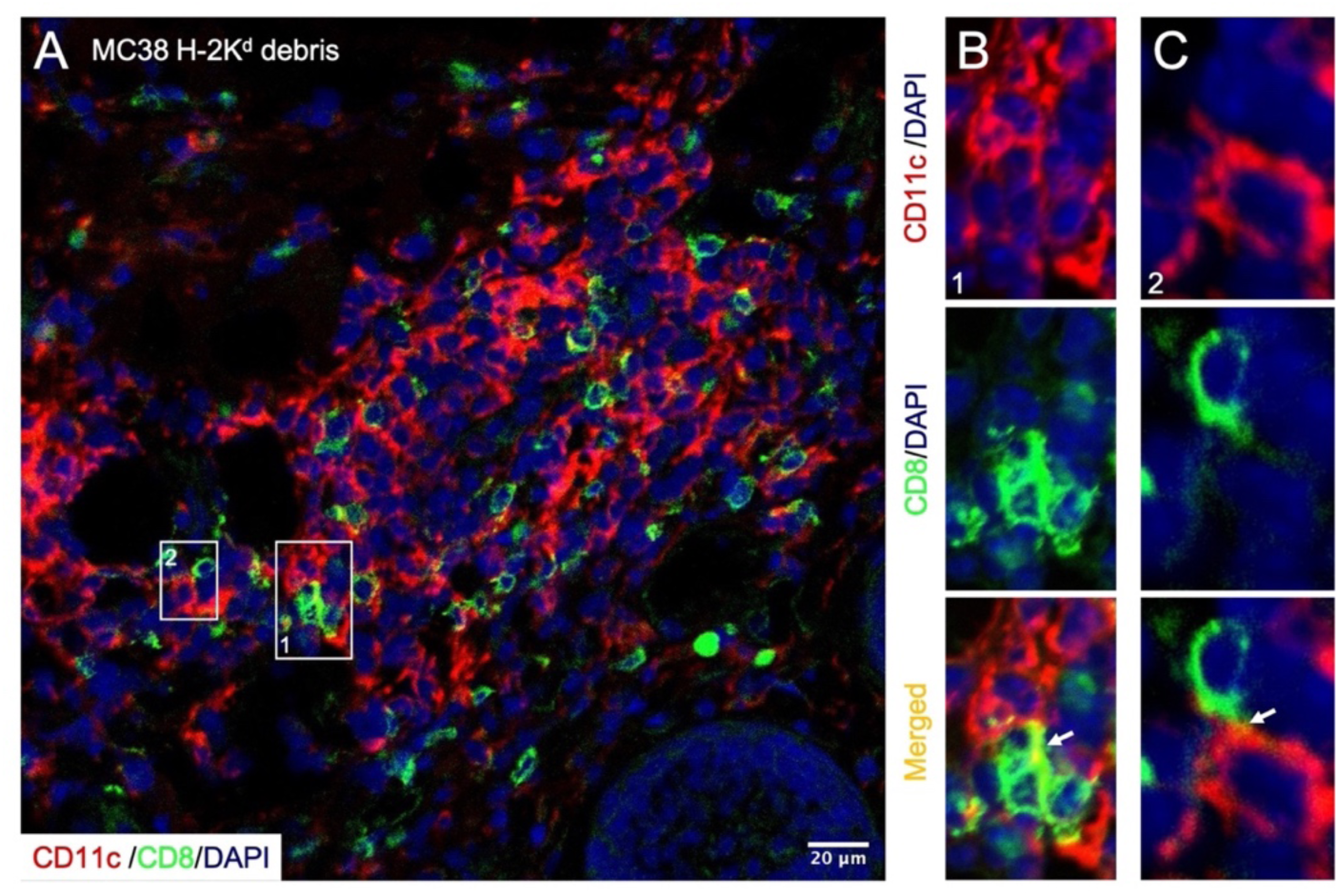
PD-1 KO mice were injected subcutaneously in the flank with Matrigel bearing debris of MC38 H-2K^d^ cells. **(A** to **C)** Representative images of plug sections at day 7 were immunostained for CD11c^+^ (dendritic cells, red) and CD8^+^ (T cells, green), and 4′,6-diamidino-2-phenylindole (DAPI) (nuclei, blue). **(A)** Visualization of host cells infiltrating Matrigel. **(B** and **C)** Higher magnification of the area inside the region of interest marked with the white box in (A), visualizing the possibility of direct cell-cell interaction between CD11c^+^ cells and CD8^+^ cells as indicated by white arrows.

## 4. Discussion

During the past few years, immune checkpoint blockade has provided remarkable outcomes to unleash the immune system against tumors. However, the majority of patients were unable to receive benefits from the PD-1 therapy (9), and some patients acquired resistance after a period of the initial response, leading to disease recurrence (74). Thus, it is indispensable to improve the success rate of PD-1 blockade in a wide variety of cancer. Furthermore, combination therapy has been widely accepted as one of the most striking strategies to overcome resistance to immunotherapy (23, 46–48). In this study, we have attempted to bolster responsiveness to PD-1 checkpoint inhibitors by exploiting allogeneic murine MHC-I genes.

The downregulation of MHC-I molecules is often associated with tumor immune evasion (27, 28). Intriguingly, the presence of tumor antigen-bound MHC is evidently important to improve the efficacy of PD-1 therapy (75). Theoretically, the introduction of allogeneic MHC-I-overexpressing vectors might improve the MHC-I-restricted immune responses mediated by cytotoxic T cells (*i.e.*, direct alloantigen recognition). The expression of MHC molecules may further help to direct immune cells to the site of antigen presentation. On the other hand, to initiate an immune response, antigen-presenting cells (APCs) such as dendritic cells must display antigens through MHC. Therefore, the uptake of allogeneic MHC by APC in the host cells may help to accelerate immune response (*i.e.*, indirect alloantigen recognition). Most notably, the presentation of allogeneic MHC-I by APCs is extremely important to aid the recruitment of immune cells to the tumor sites.

Paradoxically, although both cancer cell lines used in this study expressed high amounts of endogenous H-2K^b^ molecules, however, the parental tumors were not rejected in tested animals. This finding emphasizes that there is a threshold required for the induction of effective anti-tumoral immunity. It has been reported that even cancer cells that are capable of efficiently expressing their neoantigens through class I MHC still failed to be recognized and killed by T cells specific for this particular tumor, pointing out the presence of non-functional tumor antigens (76) and immune escape mechanism.

While the T lymphocyte response elicited by progressive tumors is typically insufficient to achieve tumor rejection, defects have been described in several stages of the immune response in the tumor-bearing host including failure in antigen-presenting machinery (26, 27, 31, 34). Transfer of MHC-I genes into tumor cells could lead to enhancement of antigen presentation and tumor elimination. We thus attempted to evaluate the therapeutic effect of allogeneic MHC-expressing tumors *in vivo*. Satisfactory results were achieved by the MC38 H-2K^d^ and MCA-205 H-2K^d^ tumor models. MC38 carcinomas were shown to be partially resistant to the chronic absence of PD-1 (Fig. 5). MC38 H-2K^d^-modified cancers showed full sensitivity to rejection by the immune system of both naïve PD-1 KO and WT mice, represented by spontaneous tumor regression (Fig. 7). In the MCA-205 tumor model which conferred high responsiveness to PD-1 deficiency, the challenged mice were completely benefited by H-2K^d^–mediated anti-tumor immunity, even without any blockade of PD-1/PD-L1 pathway (in WT mice). Notably, the expression of cell surface H-2K^d^ successfully converted non-responders into responders.

Furthermore, in the context of concomitant tumor immunity, PD-1 KO mice were prophylactically prepared for the challenge of unresponsive parental cancer cells by administering the highly immunogenic MC38 H-2K^d^ inoculums. Intriguingly, all PD-1 KO mice gained immunity against the aggressive MC38 tumor and became tumor-free (Fig. 11). This finding leads to another open question regarding in what kind of fashion the effector immune cells could recognize the allogeneic H-2K^d^ molecules. To test this, the sensitization experiment was elaborated into distinct approaches in which the growth-arrested and dead MC38 H-2K^d^ tumors were employed. The compromised MC38 H-2K^d^ tumors never grew in PD-1 KO mice, validating the efficacy of MMC and freeze-thaw treatment. More importantly, regardless of the cancer cells treatments prior to inoculation, we found that all tested PD-1 KO animals became immune to the challenge of original MC38 cells in the distant site (Fig. 12). Interestingly, the successful sensitization was proven to be reliant on PD-1 deficiency and tumor-specific immunity (Fig. 16, Fig. 17, and Fig. 19).

Consistently, we have demonstrated that the sensitization regimen elicits an efficient tumor rejection and persistent, long-lasting immunity. Specifically, the debris of MC38 H-2K^d^ cells was capable of providing durable anti-tumor immunity (Fig. 14B and 14C). This unanticipated yet surprising finding of optimum anti-tumor immune response mediated by allogeneic H-2K^d^-containing cell debris has sparked a potential provision for immunogenic cell death. Intriguingly, this result was accomplished exclusively in the context of PD-1 deficiency in combination with allogeneic H-2K^d^, whereas similar treatment repeated in WT animals, or by employing MC38 PT tumors, were crestfallen (Table 1).

**Table 1.** Induction of tumor immunity against MC38 cells.

| Mouse | Cells used for sensitization<br>(primary tumor) |  | Rejection of<br>challenged<br>MC38 PT cells | Long-term<br>immunity against<br>MC38 PT cells |
| --- | --- | --- | --- | --- |
| PD-1 KO | MC38 PT | Live cells | Yes* | N/A |
|  |  | Growth-arrested cells | Yes | N/D |
|  |  | Cell debris | **No** | N/A |
| PD-1 KO | MC38 H-2K<sup>d</sup> | Live cells | Yes | +++ |
|  |  | Growth-arrested cells | Yes | ++ |
|  |  | Cell debris | **Yes** | ++++ |
| WT | MC38 H-2K<sup>d</sup> | Live cells | No | N/A |
|  |  | Growth-arrested cells | No | N/A |
|  |  | Cell debris | No | N/A |

\*Deaths due to progressive growth of primary tumor (MC38 PT)
N/A: not applicable, N/D: not determined

Rationally, findings in this study raised a possibility for the potential application of H-2K^d^ cell debris for immunization, especially in combination with PD-1 blockade therapy. A major advantage of using cell debris is its safety and feasibility as a tumor-targeting approach to provide accessible immunogenic materials. Generally, there is no established common consent for how tumors from patients should be prepared as a cancer vaccine. The long interval time between the tedious processing of tumors until they are available as cancer vaccine might eventually result in decrement of vaccine efficacy as the progressively growing tumors accumulated a much higher burden (77). Allogeneic cancer vaccines may then serve as potent prophylaxis, a readily available option to administer before tumors reach a more advanced stage and become overwhelming.

Moreover, the strong immunity induced by allogeneic H-2K^d^ molecules in combination with global PD-1 absense leads to spontaneous tumor rejection in the MC38 cancer cell model. This finding showcases that antitumor immune responses were activated before tumors were completely rejected. Consequently, characterization of the early immune cell infiltration becomes the next interesting point to investigate. A clever strategy to study the immunological events at the nascent tumor site was achieved by embedding the inoculated tumor cells in Matrigel. Matrigel is a soluble basement membrane that is extracted from Engelbreth-Holm-Swarm (EHS) mouse sarcoma, a tumor rich in extracellular matrix membrane proteins (laminin, collagen IV, heparan sulfate, proteoglycan, and entactin) (78). Accordingly, Matrigel represents a natural tumor microenvironment and is also permissive for host cells penetration.

In this study, the Matrigel plugs containing live MC38 H-2K^d^ cells were heavily infiltrated by CD11c^+^ and CD8^+^ lymphocytes. Several possibilities of intratumoral cell-cell interaction between CD11c^+^ and CD8^+^ T cells were particularly found in both live and dead MC38 H-2K^d^-containing Matrigel plugs (Fig. 21 and Fig. 22). Tumor-associated DCs are assumed to endocytose cellular debris and transport tumor antigens to the draining lymph node where they are primed with T cells and induce T cell activation (79–82). Furthermore, cross-talk between DCs and CD8 T cells in the tumor niche and tumor-draining lymph nodes has been associated with improved efficacy of tumor-killing activity (83–85). Although this study does not rule out that other cell subsets expressing CD11c and CD8 might exist, the initial attempt by immunostaining suggested that the injection sites were massively infiltrated by cell types associated with APCs and tumor-killing T cells.

Most excitingly, our results provide compelling evidence for the potential augmentation of antitumor immune response mediated by allogeneic mouse MHC class I. Prior to the H-2K^d^ gene introduction, cancer cells used in this study were growing robustly, leading to zero survival rate of tumor-bearing mice. By appreciating the fact that the presence of H-2K^d^ alone could positively reverse the outcomes of tumor progression suggested that this approach appears to be effective. However, some limitations are worth noting. The unproved tenet of cancer treatment is that the earlier the tumor is found, particularly at the lower burden, the more likely the cancer immunotherapy is to be successful. Sensitization with the allogeneic molecule in this study preferably leaning towards the prophylactic setting. More in-depth studies will be needed to evaluate the therapeutic potential of H-2K^d^ cell debris in combination with anti-PD-1 antibody-mediated checkpoint blockade. Future works should therefore investigate the intricate roles of host-immune cells in recognizing and choreographing the H-2K^d^-mediated immune response. Nevertheless, the empirically grounded findings in this study reveal an exciting development to enhance the benefit of PD-1 therapy.

## Acknowledgments

We thank Tasuku Honjo (Kyoto University) for providing us with the PD-1 KO mice. We also thank Takashi Akazawa, Yu Mizote, and Hideaki Tahara (Osaka International Cancer Institute) for providing us with the MC38 and MCA-205 cell lines and with various kinds of technical/intellectual supports in the field of tumor immunology. This study was supported by JSPS KAKENHI Grant-in-Aid for Scientific Research B (21H02717) and for Challenging Research (Exploratory) (22K19508); ONO Pharmaceutical Co., LTD.

## Author contributions

K.A. Paramitasari performed all the experiments based on her own ideas, analyzed the data, and wrote the manuscript. Y. Ishida conceived the project, acquired the research grants, supervised the research, and wrote the manuscript.

## Conflicts of Interest

Y.I. received a research grant from ONO Pharmaceutical Co., LTD. (Osaka, Japan). The other author declares no conflicts of interest.

## Notes

### Competing Interest Statement

Y. Ishida received a research grant from ONO Pharmaceutical Co., LTD. (Osaka, Japan). The other author declares no conflicts of interest.

